# Gut dysbiosis was inevitable, but tolerance was not: temporal responses of the murine microbiota that maintain its capacity for butyrate production correlate with sustained antinociception to chronic morphine

**DOI:** 10.1101/2024.04.15.589671

**Authors:** Izabella Sall, Randi Foxall, Lindsey Felth, Soren Maret, Zachary Rosa, Anirudh Gaur, Jennifer Calawa, Nadia Pavlik, Jennifer L. Whistler, Cheryl A. Whistler

## Abstract

The therapeutic benefits of opioids are compromised by the development of analgesic tolerance, which necessitates higher dosing for pain management thereby increasing the liability for drug dependence and addiction. Rodent models indicate opposing roles of the gut microbiota in tolerance: morphine-induced gut dysbiosis exacerbates tolerance, whereas probiotics ameliorate tolerance. Not all individuals develop tolerance which could be influenced by differences in microbiota, and yet no study design has capitalized upon this natural variation. We leveraged natural behavioral variation in a murine model of voluntary oral morphine self-administration to elucidate the mechanisms by which microbiota influences tolerance. Although all mice shared similar morphine-driven microbiota changes that largely masked informative associations with variability in tolerance, our high-resolution temporal analyses revealed a divergence in the progression of dysbiosis that best explained sustained antinociception. Mice that did not develop tolerance maintained a higher capacity for production of the short-chain fatty acid (SCFA) butyrate known to bolster intestinal barriers and promote neuronal homeostasis. Both fecal microbial transplantation (FMT) from donor mice that did not develop tolerance and dietary butyrate supplementation significantly reduced the development of tolerance independently of suppression of systemic inflammation. These findings could inform immediate therapies to extend the analgesic efficacy of opioids.

## INTRODUCTION

A surge in research has demonstrated the importance of the gut microbiota to neurological function, leading to the recognition of a gut-microbiota-brain (GMB) axis ^1–5^. Whereas the brain influences the gut microbiota by control of gut motility and thereby transit, the microbiota signals the brain via production or modification of neurotransmitters, hormones, microbial byproducts, and neuroactive metabolites^4, 6, 7^. The gut microbiota is often referred to nondescriptly as commensals, but many are mutualistic symbionts that support host health. Some others are pathobionts, which are correlated with, or capable of causing, disease in a context-specific manner. The high load of pro-inflammatory microbial antigens at the microbiota-intestinal mucosa interface, produced even by mutualists, requires that the host immune responses be trained by the microbiota to be tolerogenic and not over-react, but to still appropriately clear commensals that breach the gut barrier^8^. Paramount among neuroactive signals in the GMB-axis produced by the gut microbiota that maintain homeostasis - balance of gut and host - are various short-chain fatty acids (SCFAs) generated during fermentation of dietary fiber. SCFAs can exert direct local effects by inhibiting host histone deacetylases, thereby epigenetically modifying mucosal immunity to attenuate inflammation, while enhancing gut barriers^9–13^. In addition, as ligands of host G protein-coupled receptors (GPCRs), SCFAs regulate key functions of the gut, including transit and mucus secretion ^6, 9–13^. Upon entering the circulatory system, SCFAs can also signal the brain and other organ axes ^6, 9, 13, 14^. SCFAs that reach the central nervous system cross and bolster integrity of the blood brain barrier and promote neurogenesis and neuroplasticity^9^. Disruption of GMB signaling cascades contributes to cardio-metabolic and gastric system diseases linked to inflammation and a plethora of neurobiological dysfunctions, including opioid use disorder (OUD)^15–22^.

The utility of opioids for treating chronic pain is compromised by the development of tolerance to its analgesic effects, which necessitates dose escalation that can produce physical dependence and place an individual at heightened risk for the development of an OUD. Opioids mediate their effects through activation of G_i_-coupled GPCRs, including the primary target of opioid analgesic drugs, the mu opioid receptor (MOR), which is expressed in the central and peripheral nervous systems, including the enteric nervous system^23^. MORs are also expressed by immune cells, some of which reside in the lamina propria of intestinal mucosa^24^. Many mechanisms have been hypothesized to contribute to analgesic tolerance. Broadly, these fall into two classes: loss of receptor function (receptor desensitization), and homeostatic cellular adaptations that recalibrate baseline signal transduction thereby masking signaling from functional MORs^25^. The processes by which distinct types of MOR-expressing cells adapt to counteract repeated MOR stimulation by chronic opioid likely differ by cell type^25, 26^. Crucially, in MOR-expressing cells such as neurons that transmit sensation, known as nociceptors, the adaptations to chronic opioid drug cause a loss of morphine analgesia, or antinociceptive tolerance to drug^27^. Removal of drug can produce rebound withdrawal effects that include a heightened sensation of pain known as hyperalgesia^28^, and severe gastric pain^29^. These rebound effects indicate that, in at least some tolerant states, MORs are present and functioning and that removal of drug unmasks the homeostatic cellular adaptations that produced the tolerance^30^. In humans and even in genetically inbred mice exposed to the same dose of opioid, the degree of antinociceptive tolerance varies, indicating factors beyond genetics or degree of drug exposure contribute to loss of opioid potency^31–33^. Understanding what underlies this variability in response to chronic opioid could produce new mechanistic insight. Preventing or delaying analgesic tolerance would reduce the need for dose escalation, thereby mitigating the risk of overdose and OUD development, and improving the utility of opioids^34^. Through their activation of MORs in the gastrointestinal tract, opioids impair motility and electrolyte secretion leading to constipation^35, 36^ and alter the gut microbiota^10, 37–42^. Gut microbiota dysbiosis is implicated in opioid-induced changes in drug reward^42, 43^, dependence^39, 44, 45^, hyperalgesia^42, 46^, and analgesic tolerance^17, 37, 40^. Several studies document that morphine-induced gut microbiota dysbiosis and subsequent translocation of pathobionts across a morphine-compromised gut barrier trigger systemic inflammation that exacerbates the development of tolerance^26, 34, 40, 47^. In agreement with this model, eliminating gut commensals prevents tolerance, and introduction of a common probiotic harboring *Lactobacillus* and *Bifidobacteria* or replenishment of gut commensals via fecal microbial transplantation (FMT) from opioid-naïve donors counters dysbiosis and antinociceptive tolerance^37, 40, 47, 48^. Importantly, whereas opioid-driven dysbiosis contributes to tolerance, it is not sufficient, as FMT from opioid-induced tolerant donors does not compromise morphine antinociception^40, 48^ though it can accelerate opioid-driven tolerance^40^. Indeed, opioid stimulation of MOR is necessary for tolerance as demonstrated by the ability of a peripherally restricted MOR antagonist to counter tolerance even in the presence of inflammation resulting from impaired barrier function^46, 47^. Whether microbiota dysbiosis has a synergistic causative effect or tangentially influences tolerance, or if inflammation is its primary mediator, is yet unknown and awaits more mechanistic studies.

Importantly, the foundational studies above all minimized and collapsed natural variation in animal and microbiome responses to opioid by using high doses of opioid and aggregating data. Here, we designed a study to leverage both natural variability in degree of antinociceptive tolerance to morphine and microbiota composition with the goal of identifying informative native microbiota signatures that could explain individual differences in the development of tolerance. We analyzed temporal changes in microbiota abundance, presence, variability, stability and interrogated the potential role of community interactions and metabolic functions^49–52^ to expand our understanding of how opioid-induced gut dysbiosis contributes to liability of tolerance. We demonstrated that morphine-induced dysbiosis is inevitable, but tolerance is not. Furthermore, since morphine-induced progressive dysbiosis occurred in all mice, this largely obscured temporal signatures differentiating tolerant from non-tolerant mice that were only discernable by comparative approaches that control for common morphine-induced changes. Whereas tolerant mice shared few distinguishing patterns other than expected higher abundance of pathobionts, mice that did not develop tolerance displayed more shared and predictive features, pointing to a protective role of the microbial neuroactive metabolite butyrate, for preventing the development of tolerance.

## MATERIALS AND METHODS

### Animal Use and Care

Male wildtype C57/BL6J mice, *Mus musculus* (Jackson Laboratory) age matched were used for both the oral morphine paradigm (n=16, 8-week-old upon entering lever training with 0.2% saccharine reward, and 11-week old at first morphine exposure; Figure 1^53^), and subcutaneous (s.c.) morphine paradigm (n=52, 8-week-old at start of dietetic pre-treatment and 10-11-weeks at morphine exposure; Figure 7). An additional cohort of 8 (n=4 female and n=4 male, 11-week at first morphine exposure) wild-type C57/BL6J mice bred in-house at UC Davis were also subjected to the oral morphine paradigm and used as a test dataset for assessing the specificity and accuracy of microbiota associations in prediction models of morphine exposure. All mice had food *ad libitum* and were provided running wheels for enrichment. Mice in the oral morphine paradigm had two bottles, one with water and one with morphine, and they were individually housed allowing for the measurement of morphine consumption. Mice in the s.c. paradigm were communally housed (3-4 mice per cage) to minimize discomfort since all mice received the same morphine dosing. Mice used for assessment of dietetic supplementation on tolerance had access only to water supplemented with either sodium butyrate (Sigma-Aldrich) or pH-matched (9.30 - 9.50) monosodium citrate (EMD) as a control at 100 mM, a concentration informed by a survey of prior studies evaluating benefits of butyrate^54, 55^. Bottles containing freshly prepared and filter-sterilized butyrate and citrate were changed every three days for 3-weeks prior morphine exposure and for the duration of the experiment^48, 54, 56^. All procedures involving animals were reviewed and approved by the Institutional Animal Care and Use Committees of the University of California, Davis (IACUC protocol number: 22085) or University of New Hampshire (IACUC protocol number: 230101).

**Figure 1:**
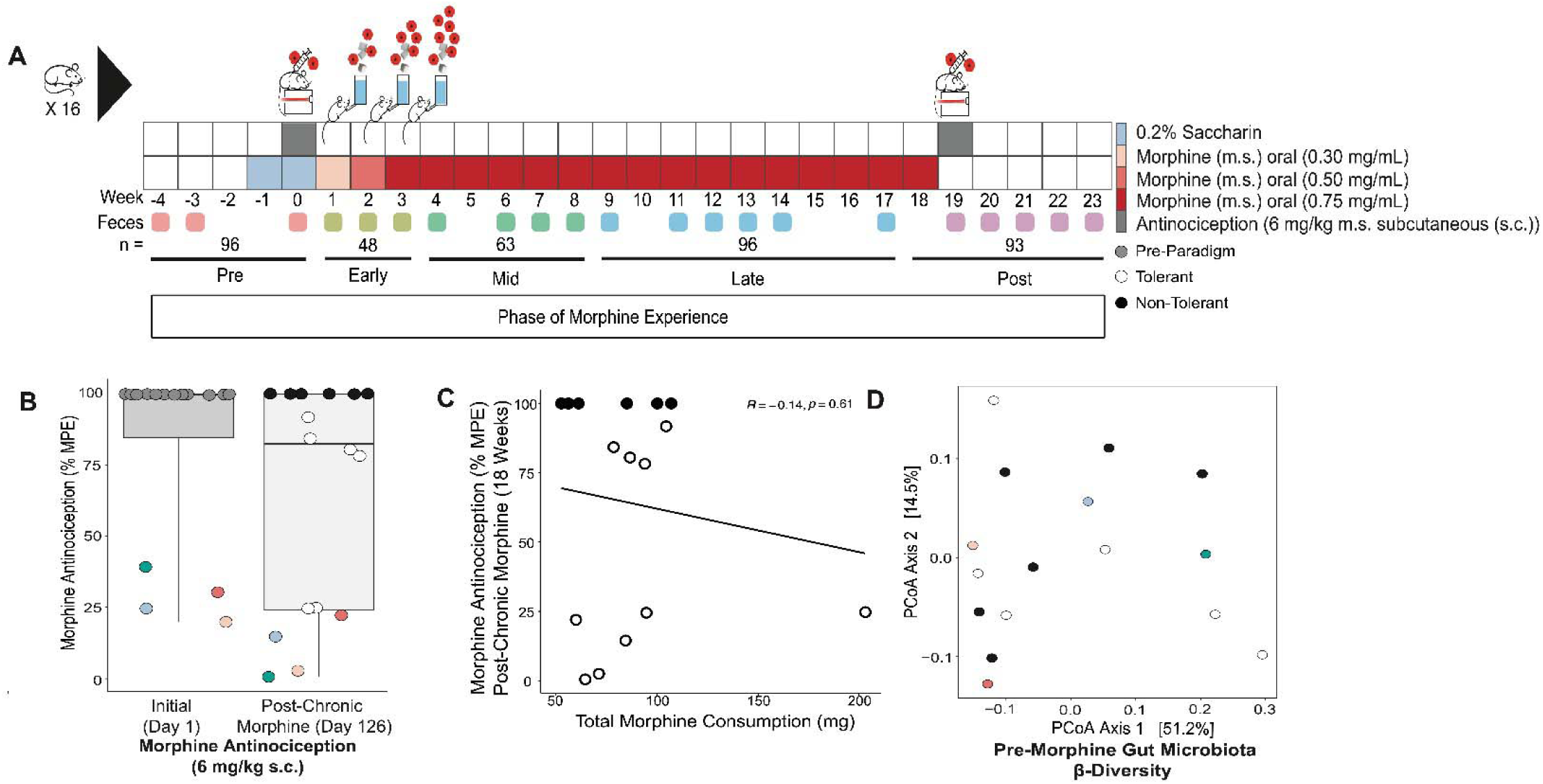
Natural variability in antinociceptive tolerance to chronic morphine does not correlate with the amount of morphine mice voluntarily consumed. A) Mouse (n=16) chronic morphine oral self-administration paradigm where voluntary morphine consumption was monitored ^53^. Fecess were collected and used to generate 16S (V4-V5 region, see methods) for microbiota analysis. B) Morphine antinociception (6 mg/kg s.c., ED_80_) was determined pre- and post-chronic morphine (week 19) using a radiant heat tail flick assay to determine tolerance to morphine. Data is reported as maximum possible effect (% MPE). Colored individuals exhibited low starting antinociception which decreased further after chronic morphine consumption consistent with tolerance C). Linear regression of total morphine consumption with degree of tolerance (y = 78 - 0.16x; R2 = −0.14; Pearson’s correlation p-value = 0.61). D) Pre-morphine gut microbiota β-diversity (Bray-Curtis dissimilarity) was visualized by principal coordinates analysis (PCoA). See Supplemental Table 1 for details.

### Morphine and Naloxone Administration

For the 18-week oral self-administration paradigm (n=16; ^53^ Figure 1, Figure 2C), mice had access *ad libitum* 5 days per week to both water and water supplemented with morphine sulfate (Mallinckrodt Pharmaceuticals, St. Louis, MO) sweetened with 0.2% saccharin to improve palatability, where the concentration of morphine sulfate was gradually escalated from 0.3 mg/mL, to 0.5 mg/mL, to a final concentration of 0.75 mg/mL during the first three weeks. Each week, mice then had 2 days where only water was available. Voluntary morphine consumption was determined from bottle weight. Morphine treatment mice in the non-contingent subcutaneous (s.c.) morphine paradigm received either 8 mg/kg morphine (Cayman Chemical, Ann Arbor, Michigan) in sterile 0.9% saline (Patterson Veterinary Supply, Greeley, CO) once each day for 4 days (n=14) or 10 mg/kg of s.c. morphine once each day for 9 days (n=78). Immediately following some assays for nociception using the ED_90_ dose of morphine (5 + 2 mg/kg) on day 10 of the s.c. paradigm, mice were treated with 5 mg/kg s.c. naloxone, an antagonist of MOR, to displace morphine allowing observation of reversal of morphine-impaired constipation, where the number of fecal pellets produced in 20 minutes was recorded for each mouse, and where a greater number of pellets represents morphine-driven decreased transit.

**Figure 2:**
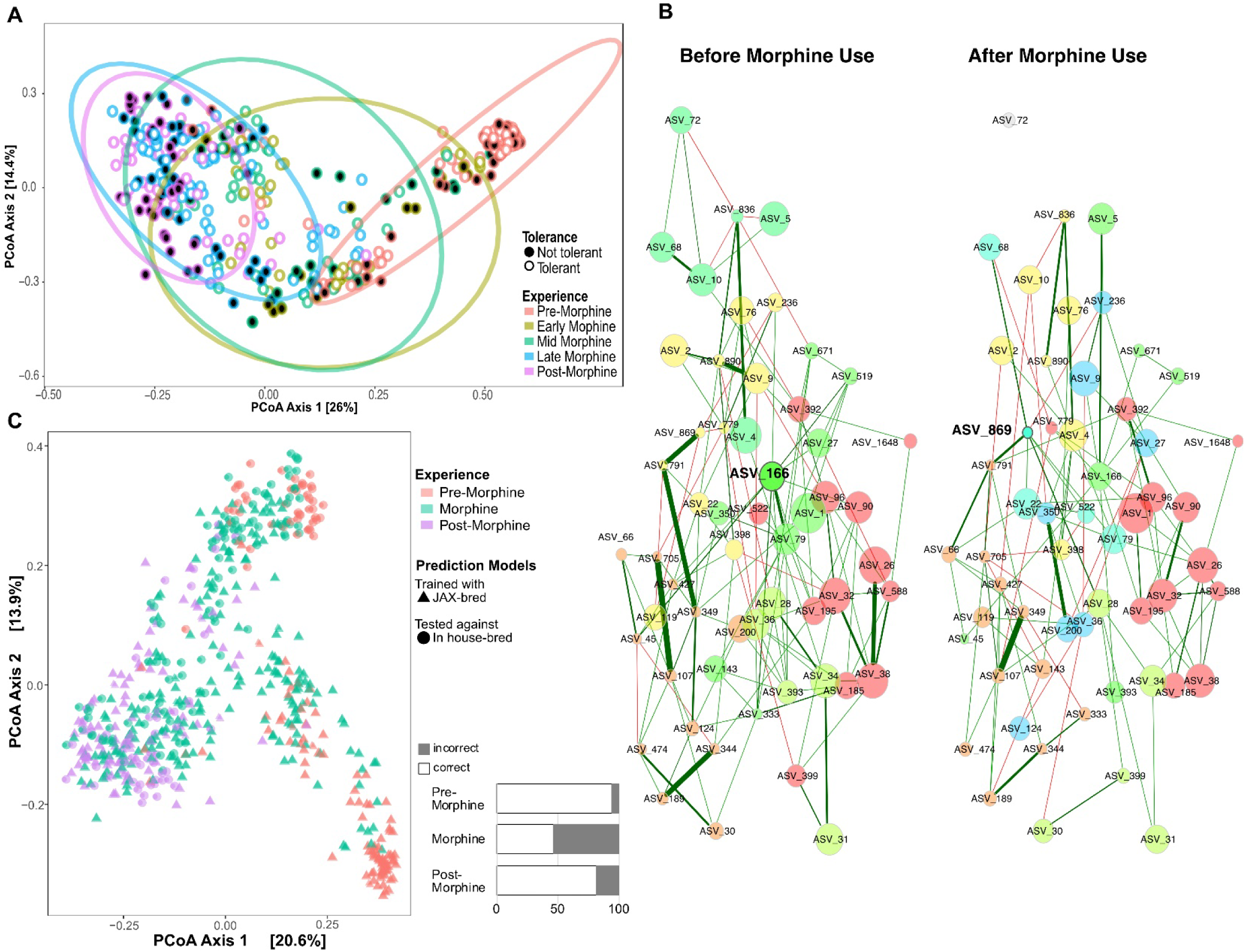
Chronic morphine induced predictive dysbiosis in all mice. A) Temporal changes in gut microbiota β-diversity (All ASVs, Bray-Curtis dissimilarity) as visualized by principal coordinates analysis (PCoA). See Supplemental Table 1 for details. B) Comparison of bacterial association networks before (left) and after 18 weeks (right) of oral morphine using the SPRING method in NetCoMi^94^. Edges are colored by sign (positive: green, negative: red), representing estimated associations between community members. *Adlercreutzia* (ASV_166) and *Streptococcus* (ASV_869) are the only two community members identified as hubs (bolded text and borders). Eigenvector centrality was used for scaling node sizes and colors representing clusters of genera (determined using greedy modularity optimization). Clusters have the same color in both networks if they share at least two taxa. Nodes that are not connected in both groups were excluded. Corresponding taxa names detailed in Supplemental Table 3. C) Predictive modelling of morphine-exposed microbiota (using R program indicspecies)^90^. β-diversity (All ASVs, Bray-Curtis dissimilarity) as visualized by principal coordinates analysis (PCoA). The starting composition of microbiota of mice that trained the models (JAX-bred, orange triangles), which group in the lower right corner, and of mice that tested the models (in-house-bred, orange circles), which group in the upper right corner, differ but converge upon exposure to morphine. The right bar graphs show % accuracy of the models to predict whether microbiota from in-house-bred mice resulted from morphine exposure (as % correct/incorrect). See Supplemental Tables 4 and 5 for details.

### Determination of Nociception and Morphine Antinociception

The radiant heat tail flick assay (Tail-flick Meter, Columbus Instruments) was used to assess baseline nociception and morphine antinociception. Light intensity was set to produce a baseline tail-flick latency of ∼2.0 seconds with a maximum cutoff of 6 seconds (for oral paradigm) or 5 seconds (for s.c. paradigm) to minimize tissue damage. Latency was measured without morphine, and then 30 min after morphine injection. For the oral paradigm, mice received 6 mg/kg morphine (ED_80_) ^57^ before the study’s start and again at the study’s completion (Figure 2A). For the s.c. paradigm with FMT or butyrate/citrate feeding, antinociception was determined on days 1, 7 and 10 at 30 minutes after a 5 mg/kg dose of morphine (ED_60_), followed by a second dose of + 2 mg/kg morphine (combined ED_90_) and a second latency was measured 30 minutes later. Saline control mice (n=16) only received morphine for the antinociception tests on day 10. The percent maximum possible effect (%MPE) was calculated with the following formula: 100 * [(drug response time-population baseline latency) / (cutoff time-population baseline latency)]. All tail flick tests were blinded to prevent bias. Dose– responses were evaluated with 12-week-old male mice to optimize doses for maximum statistical power and effect size using power analysis. These and the cumulative dose responses for evaluating tolerance were fit using GraphPad Prism. The ED_50_ values of morphine and their 95% confidence intervals were calculated and a shift in ED_50_ from day 1 to day 10 was calculated. Statistical support for a change in morphine potency was determined by a two-tailed P test, testing the null hypothesis that the dose response slope on day 10 did not differ from the slope on day 1. Mice with the same %MPE at both doses (either at ceiling at 5mg/kg, or at floor at 5+2mg/kg) were not used to calculate change in slope. Mice with sustained antinociception at the end of either the oral or s.c. paradigm (100% MPE) were classified as “non-tolerant”, whereas mice with reduced morphine antinociception from day 1 values (<100% MPE) were classified as “tolerant”.

### Fecal Microbiota Transplantation

To generate feces for fecal microbial transplantation (FMT), a small cohort of male wildtype C57BL/6J mice (n = 14) received 8 mg/kg s.c. morphine for 4 days were assessed for tolerance on day 5 with 6 mg/kg s.c. morphine using the radiant heat tail flick assay. A total of 2.7 g of feces pooled from multiple individuals of two populations, one that did not become tolerant (maintained 100% MPE) and one that did become e tolerant (displayed < 100% MPE), was collected and immediately cryopreserved at −80 °C both for use in FMT and for microbiota analysis. These feces were sufficient for inoculation of each treatment group (n = 14 for each FMT treatment group; n = 9 receiving s.c morphine and n = 5 receiving s.c. saline carrier). The feces from tolerant and non-tolerant mice were subsequently defrosted on ice and suspended in a total of 32.4 mL of cold (4°C) phosphate-buffered saline (PBS) containing 10% glycerol^48^. The suspension was homogenized and then centrifuged at 800g for 3 minutes ^48^ and the supernatant of each was aliquoted in 2.8 mL volumes in separate tubes and cryopreserved at −80 °C. Additionally, 100 µL aliquots of each homogenized suspension from pooled tolerant and pooled non-tolerant mice were reserved for microbiota (16S) analysis. Upon entering the FMT study, recipient mice were administered 100 µL of the fecal supernatant twice daily ^48^ for 9 days via intragastric gavage: Feces were collected from all mice on day 0 (pre-treatment) and 10, and stored at −80 °C. Analyses of the microbiota of tolerant and non-tolerant pooled donor feces were limited due to lack of statistical power (of only two samples) but comparison of compositional differences were illustrated using the ps_venn function in the MicEco R package (v0.9.15) ^58^ to determine the overall number of unique and shared amplicon sequence variants (ASVs).

### Statistical Analysis of Animal Data

Linear regression analyses using Pearson’s correlation coefficient^59^ determined whether morphine antinociception significantly correlated with total morphine consumption using the R package ggpubr (v0.6.0)^60^. Normality of data was assessed using a Shapiro-Wilk test and homogeneity of variance was assessed using Levene’s test. Data with a p-value greater than 0.05 was considered normal. One-way ANOVA followed by post-hoc analysis using Tukey’s Test for multiple comparisons was used to compare differences between groups where assumptions for normality and homogeneity of variance were met. The Kruskal-Wallis test followed by pairwise Wilcox test with corrections for multiple comparisons was used when assumptions of normality and homogeneity of variance were not met. All statistics were conducted using the R package stats (v4.3.0)^61^. Visuals were generated using R 4.2.0 software, GraphPad Prism, BioRender, and some composite figures were modified with Adobe Illustrator.

### 16S rDNA Library Preparation and Sequencing

Individual mice were placed in clean cages with no bedding for an hour to defecate normally. Fresh feces were collected and immediately cryopreserved at −80 °C. Samples were randomized across 96-well plates using the R package wpm (v1.14.0) to minimize introduction of batch effects from multiple DNA extractions and sequencing runs^62^. Feces were lysed using the Qiagen TissueLyser II (Qiagen, Germantown, MD) for 5 minutes at 30 Hertz. Microbial genomic DNA was extracted from frozen feces (∼100 mg, or 5 pellets) with the ZymoBIOMICS 96 MagBead DNA Kit with lysis tubes (Zymo Research, Irvine, CA), according to manufacturer’s protocol. DNA was quantified using the iT dsDNA Broad-Range kit (Invitrogen, Life Technologies, Grand Island, NY) using a Tecan M200 plate reader (excitation 480nm/emission 530nm) (Tecan, Männedorf, Switzerland) in a Costar black, clear bottom plate (Corning, Corning, NY). Genomic DNA samples were standardized (11 ng/µL) and freshly diluted to 1.1 ng/µL prior to two successive polymerase chain reaction (PCR) amplifications. The V4-V5 16S variable regions were first amplified in 10 µL reactions in triplicate using the NEB Q5 Hot Start HiFi 2X master mix (New England BioLabs, Ipswich, MA) and 515F and 926R primers as described^63, 64^, and modified to include the forward and reverse primer pad and linker at the 5’ end of each primer. The cycling parameters were as follows: 98°C for 2 min, followed by 22 cycles of 98°C for 10 s, 53°C for 15 s, and 72°C for 15 s, and completing with 72°C for 2 min. Following estimation of size, quality, and quantity of amplicons by gel electrophoresis, pooled amplicons were enzymatically cleaned with ExoSap-IT (Applied Biosystems, Thermo Fisher Scientific, Waltam, MA) and diluted 1:5 in nuclease-free water prior to indexing. In the second step, amplicons were indexed using xGen UDI Primer Pairs (10 mM) in 12 µL reactions using the NEB Q5 Hot Start HiFi 2X master mix (New England BioLabs, Ipswich, MA) with the following parameters: 98°C for 30 s, followed by 10 cycles of 98°C for 10 s, 65°C for 15 s, and 72°C for 20 s, and a final step of 72°C for 2 min. Amplicon band size, quality, and quantity was verified using 2% agarose by gel electrophoresis. Indexed amplicons were pooled in a final library and purified with the Mag-Bind Total Pure NGS kit (Omega Bio-Tek, Norcross, GA). The pooled library was quantified using the Qubit dsDNA High Sensitivity kit (Invitrogen, Life Technologies, Grand Island, NY) and the Qubit 2.0 fluorometer (Invitrogen, Life Technologies, Grand Island, NY). Final amplicon libraries were sequenced and demultiplexed on the Illumina NovaSeq 6000 platform (2 x 250 bp paired end) at the Hubbard Center for Genome Studies at the University of New Hampshire (Durham, NH). Computations were performed on Premise, a central, shared HPC cluster at the University of New Hampshire (Durham, NH) supported by the Research Computing Center and PIs who have contributed compute nodes.

### Processing of 16S rDNA Sequencing Reads

All necessary components needed to generate microbial community profiles (e.g., taxonomic assignments, sequence counts, sample metadata, phylogenetic tree;) and to conduct downstream analyses were stored in an experiment-level object using the R package phyloseq (v1.40.0)^65^. For a summary of 16S processing and analysis, see Supplemental Figure 1.

Amplicon sequencing primers were removed from demultiplexed sequencing reads using the cutadapt plugin ^66^ in the QIIME 2 2020.2 pipeline^67^ before being processed using the R package DADA2 (v1.18) quality control pipeline^68^ and custom scripts for post-processing artifact removal. Taxonomy was assigned using the GreenGenes reference database (v13.8)^69^. Species, where applicable, were identified using rRNA/ITS databases with the nucleotide basic local alignment search tool (blastn)^70^. A taxonomic filter was applied to remove unclassified phyla, chloroplasts, and mitochondria, along with an additional abundance filter to remove singletons and doubletons from the dataset (phyloseq v1.40.0)^65^. Maximum likelihood phylogenetic trees with bootstraps were estimated from MAFFT (v7.305b)^71, 72^ alignments using RAxML (v8.2.10)^73^, followed by tip-agglomeration (h = 0.5) to combine similar amplicon sequence variants (ASVs) into one representative taxon sequence or taxa agglomeration to the genus level using phyloseq (v1.40.0)^65^. Sequencing counts were transformed to relative abundances (phyloseq v1.40.0)^65^ or normalized to center-log ratios (R package microbiome v1.18.0)^74^.

The presence of batch effects that may introduce bias were assessed by comparing sequenced controls using the R package MMUPHin (v1.10.0)^75^ and principal coordinates analyses (PCoA). The controls used to assess bias included 1) a ZymoBIOMICS Microbial Community Standard II (Log Distribution) to detect any bias introduced during DNA extraction, 2) a ZymoBIOMICS Microbial Community DNA Standard II (Log Distribution) to detect any bias introduced during amplification, 3) a randomized representative mouse fecal sample to detect any bias introduced from multiple sequencing runs, and 4) a no template mock DNA extraction to capture any potential contaminants.

### Analysis of Microbiota Diversity

Overall gut microbiota diversity and temporal changes assessed by weighted UniFrac^76^, unweighted UniFrac^77^ (membership), and Bray-Curtis dissimilarity (relative abundance) was analyzed using PCoA. Permutation Multivariate Analysis of Variance (PERMANOVA) tests^78^ determined any statistical differences between experiences and the development of antinociceptive tolerance throughout the paradigm using the permanova pairwise function in the R package ecole (v0.9-2021)^79^. Prior to using PERMANOVA, pairwise permutation tests^80, 81^ for homogeneity of multivariate dispersion^82–84^ determined any statistical differences in group variances of experiences and the development of antinociceptive tolerance throughout the paradigm using the betadisper and permutest functions in the R package vegan (v2.6-4)^85^.

Alpha-diversity of non-tolerant and tolerant gut microbiota were analyzed using the estimate_richness function for Shannon diversity in the R package phyloseq (v1.40.0)^65^. Average alpha-diversity and standard errors were plotted sequentially and abrupt variations over time that may represent key transitions were detected using a Bayesian analysis of change point in the R package bcp (v4.0.3)^39, 86^, which implements the Barry and Hartigan product partition model for the standard change point problem using Markov Chain Monte Carlo^87^. Timepoints with a posterior probability of change in the mean abundance greater than 0.70 of community members in tolerant or non-tolerant mice were marked on time series plots.

### Differential Abundance and Biomarker Analyses

Community members, grouped at the genus level, that were different in abundance between tolerant and non-tolerant mice, and between pre-, during, and post-morphine exposure phases of the paradigm, were determined using the differential test within the R package corncob (v0.3.1)^88^. Non-normalized counts were used with the Wald (abundance) or LRT (variability) setting within the differential test to distinguish community members associated with the covariate of interest controlling only for sequencing run and phase of morphine. The count abundance for each genus was fit to a beta-binomial model using the logit link functions for both the mean and overdispersion simultaneously. The null and non-null overdispersion models were specified with the same confounding variables (paradigm experience and sequencing plate) to identify only genera having differential abundances associated with the covariate of interest. The list of differentially abundant and/or variable community members produced was further analyzed using linear discrimination analyses of effect size (LEfSe Galaxy v1.0)^89^ to identify any community members that most likely explain differences between a covariate of interest. These community members were visualized using the amp_heatmap function in the R package ampvis2 (v 2.8.9) on a log2 color scale and the minimum abundance set to null.

### Indicator Species and Prediction Models

We used an indicator species analysis to determine community members unique to experiences and to provide evidence for the impacts of experience on the gut microbiota^90^. Furthermore, we tested whether there were combinations of up to 3 indicator species that could predict whether a sample was from a morphine exposed gut microbiota. Abundance data for the cohort was transformed into presence absence data and indicator value analysis with experience as the grouping (pre-morphine, morphine, post-morphine) was performed using the multipatt() function with 999 permutations in the R package indicspecies^90^. To determine which combinations of community members could best predict whether a sample was from morphine exposure we reduced the dataset to samples that were from pre-morphine or mid-late morphine and used the indictors() function with up to 3 combinations of community members, and subsequently the pruneindictors() function to assess which combinations were best predictors. Next, we tested whether the combinations which were only found in mid-late morphine samples (Probability A=1) could accurately predict which samples were most likely from morphine exposed microbiota from a separate cohort of 8 animals (n=4 female, n=4 male) that consisted of 40 pre-morphine, 102 morphine samples, and 15 post-morphine samples.

### Longitudinal Change Point Analysis

Abrupt variations over time that may represent key transitions in community members that were differentially abundant between tolerant and non-tolerant mice were detected using a Bayesian analysis of change point in the R package bcp (v4.0.3)^39, 86^, which implements the Barry and Hartigan product partition model for the standard change point problem using Markov Chain Monte Carlo^87^. Timepoints with a posterior probability of change in the mean abundance greater than 0.790 of community members in tolerant or non-tolerant mice were marked on time series plots.

### Analysis of Temporal Stability

The temporal stability of populations (All ASVs, grouped at the genus level) exposed to morphine was evaluated and visualized within the context of Taylor’s power law, which describes the ubiquitous macroecological relationships between the mean and variance of an individual taxon^91^. Two extensions of this law were used to examine whether the microbiota of tolerant and non-tolerant mice differ in variability or stability. The Type I power law extension was used to detect differences in temporal variability of tolerant and non-tolerant microbiota across all phases of morphine experience and the Type IV power law extension was used to examine differences in microbiota stability relative to starting composition at each phase of morphine experience^92^. When applying the Type IV extension, Taylor’s power law which is described by the formula V = aM^b^, where *V* is the variance, *a* is a scalar parameter, *M* is the mean, and *b* is an exponential parameter, the formula was log-transformed, ln(*V*) = *b*ln(*M*) + ln(*a*), where *b* is the slope parameter^92, 93^. The exponential/slope parameter b of the Type IV power law extension was standardized by subtracting the mean b of all individuals pre-morphine from each individual mouse’s value for b at each morphine experience to produce the change in slope. Positive values of the slope parameter *b* represent instability, whereas 0 or negative values of the slope parameter b represent stability relative to pre-morphine.

### Microbiome Association Networks

Microbial association networks were inferred from the gut microbiomes of mice before entering the self-administration of oral morphine phase in the paradigm compared to the gut microbiomes after 18-weeks of morphine administration to 1 week post administration. Community members were grouped at the genus level using the tax_glom function from phyloseq (v1.40.0)^65^. Seventy out of 165 taxa grouped at the genus level were at the highest frequency. SPRING associations were measured in the gut microbiota split by whether they were taken before or after morphine exposure and another iteration where microbiota were split based on development of antinociceptive tolerance. Microbial association networks for each group were inferred using the signed distance, nlambda, and rep.num (100) functions in the R package NetCoMi (v1.1.0)^94^. For reproducibility, microbial association network analyses used the random seed “12345”. Differential networks were inferred from SPRING association networks using fisher test adjusting for false discovery rate.

### Analysis of Predicted Functional Profiles

Functional profiles of gut microbiota from tolerant and non-tolerant mice were predicted from representative ASVs of 16S rDNA sequences using phylogenetic investigation of communities by reconstruction of unobserved states (PICRUSt 2.0) pipeline^95^. Gene content, represented by Enzyme Classification (EC) numbers, (i.e., gene family copy numbers of ASVs and abundances per sample) per ASV was predicted^96^ by aligning ASVs to reference sequences and placement into a reference phylogenetic tree^97–99^. MetaCyc functional pathways and associated abundance were inferred from EC number abundances^100^. Differential abundance testing of functional pathways between tolerant and non-tolerant mice was analyzed^101^ across three models, including ALDEx2^102^, DESeq2^103^, and edgeR^104^ using the R package ggpicrust2 (v1.7.3)^101^.

### Assessment of Butyrate Biosynthetic Capacity

To assess the ability of intestinal microbiota to synthesize butyrate we quantified two genes that encode proteins in the butyrate production pathways, including butyryl-CoA transferase (BCoAT;*bcoat or but*) and butyrate kinase (*buk*) using the primers *bcoat*-F 5’- GCIGAICATTTCACITGGAAYWSITGGCAYATG-3’ and *bcoat*-R 5’- CCTGCCTTTGCAATRTCIACRAANGC-3’^105^, along with Buk-5F1 5’- CCATGCATTAAATCAAAAAGC-3’, Buk-5F2 5’-CCATGCGTTAAACCAAAAAGC-3’, Buk-6R1 5’- AGTACCTCCACCCATGTG-3’, Buk-6R2 5’-AATACCTCCGCCCATATG-3’ and Buk-6R3 5’-AATACCGCCRCCCATATG-3’^106^ by quantitative real-time PCR (qPCR). Copy numbers were normalized to the bacterial 16S rRNA gene copy number amplified by the universal primer set 16S rRNA-F 5’-AGAGTTTGATYMTGGCTCAG-3’ and 16S rRNA-R 5’ACGGCTACCTTGTTACGACTT-3’^105^. The qPCR was performed in duplicate in a final volume of 20µl using PerfeCTa SYBR Green FastMix Low ROX (Quantabio, Beverly, MA) with fecal DNA (30 ng for BCoAT, 10 ng for *buk*, and 1 ng for 16S rRNA) and 100nM of each primer for *bcoat* and 16S rRNA, and for *buk* 800nM forward primers, and 1400nM reverse primers (400nM buk-6R1, 200nM buk-6R2 and 800nM buk-6R3). Cycling conditions for both BCoAT and 16S rRNA were as follows: 95°C for 3 minutes followed by 40 cycles at 95°C for 30s, 56°C for 30s, 72°C for 40s for BCoAT and 2min for 16s rRNA and a final extension at 72°C for 5 minutes^105^. The cycling parameters for *buk* were an initial denaturation at 95°C for 5 minutes followed by 40 cycles at 95°C for 30s, 52°C for 30s, and 71°C for 30s^107^. Student’s T-test determined significant differences between groups.

### Blood Serum Biomarkers of Inflammation and Bacterial Translocation

Serum was separated from clotted blood collected via terminal bleed by cardiac puncture using an 18-gauge needle and assayed for various indicators of systemic inflammation and bacterial translocation. Briefly, collected blood was allowed to clot for 30 minutes before a 10-minute centrifugation at 2,000 x g and 4°C. Each serum sample was aliquoted into replicate tubes and stored at −80°C, each of which was only thawed once for subsequent assays. Serum cytokines (IL-1β, IL-6, and TNF-α) were quantified using reagents and manufacturers protocols for the Bio-Plex Pro mouse cytokine Th17 Panel A kit from Bio-Rad (Hercules, CA, USA) at the IDDRC Biological Analysis Core laboratory (UC Davis, CA, USA).

To assess the extent of bacterial translocation from the intestines, serum was assayed for bacterial lipopolysaccharide (LPS), and for LPS binding protein (LBP) as a surrogate for liver response to bacterial translocation. Serum was diluted 1:100 in endotoxin free tris buffer pH 8.0 and LPS quantified in duplicate using the Pierce LAL Chromogenic Endotoxin Quantification Kit (Thermo Scientific, Waltham, MA, USA) with the low range standards and following manufacturer’s protocol. The resulting chromogenic substrate was quantified at OD_405_ nm using an Infinite M200 plate reader (Tecan, Morrisville, NC, USA). LBP was quantified from serum diluted 1:100 in dilution buffer using the Mouse LBP Elisa kit (Invitrogen, Waltham, MA, USA) following the manufacturer’s protocol. The resulting chromogenic substrate was quantified at OD_450_ nm using a Tecan Infinite M200 plate reader (Morrisville, NC, USA).

## RESULTS

### Not all mice developed antinociceptive tolerance to chronic voluntary oral morphine

To examine variables that contribute to differences in OUD liability, particularly antinociceptive tolerance to morphine, individually housed wild-type mice underwent an 18-week longitudinal paradigm of voluntary oral morphine self-administration, where they had *ad libitum* access to both morphine and water^53^. To determine the development of tolerance, we used reflexive tail-flick to radiant heat to measure nociception -the ability to sense- and antinociception -the ability of morphine to block sensation- with a non-contingent morphine dose (6 mg/kg, s.c.) at the start and end of the self-administration phase of the paradigm (Figure 1A). Mice in this paradigm displayed variability in day 0 baseline morphine antinociception and in tolerance to antinociception after 18-weeks of morphine self-administration (Figure 1B). More than a third of mice (6/16) maintained 100% of the maximum possible effect (MPE) to morphine antinociception (i.e. non-tolerant) after 18-weeks of self-administration. As a population, after 18-weeks of morphine, mice in our paradigm did not display a significant decrease in baseline nociceptive threshold in the absence of morphine (i.e. hyperalgesia), despite consuming, on average, 25 mg/kg of morphine daily (range of 5-120 mg/kg daily; Figure 1C & Supplemental Figure 2). Tolerance to morphine (e.g. decreased % MPE from individual day 0 measurements) did not correlate with a change in baseline nociceptive threshold in the absence of morphine indicating that tolerance was not driven solely by hyperalgesia (Supplemental Figure 2B). Importantly, individual variation in morphine tolerance also did not correlate with the amount of morphine each mouse consumed (Figure 1C), indicating that variables beyond degree of drug intake influenced the development of tolerance. Individual pre-morphine microbiota, as determined from 16S rRNA libraries generated from feces, was highly variable (Figure 1D, *p*_PERMANOVA_ = 0.0001, Supplemental Table 1) even though the mice were the same age, sex, inbred genotype, and from the same commercial source. This led us to hypothesize that holobiont genetic variation in the form of the hosts’ starting microbiome and/or its response to morphine might explain differences in the development of antinociceptive tolerance.

### Morphine induced progressive dysbiosis in all mice regardless of whether they did or did not develop tolerance

We capitalized upon our unique experimental design of intensive sampling to map the temporal impact of morphine on gut microbiota, thereby capturing snapshots of mouse guts at different phases of morphine experience. We then evaluated whether differences in microbiota were associated with variability in tolerance (Figure 2 and Supplemental Figure 3; details provided in Supplemental Tables 1 & 2). Preliminary exploration of microbiota using Bray-Curtis dissimilarity of all amplicon sequence variants (ASVs) demonstrated how morphine caused progressive shifts in bacterial community assemblages (*p*_PERMANOVA_ ≤0.01 for all pair-wise comparisons, Supplemental Table 1), even in mice that did not develop antinociceptive tolerance to morphine (Figure 2A, Supplemental Table 1). The microbiomes of tolerant and non-tolerant mice did not form separate clusters at any phase of the paradigm, and because of this, it is apparent that the experience of morphine was the primary driver of microbial communities (Figure 2A and Supplemental Figure 3). We also investigated whether morphine increased variability as a dimension of dysbiosis (Supplemental Figure 4). Early morphine exposure increased the number of variable genera relative to pre-morphine, and as morphine exposure continued, the number of variable genera increased again later in the paradigm and remained variable even post-morphine.

To further visualize dysbiosis, we examined how global network properties and differential networks of microbiota were impacted by morphine, as context for identifying differences between tolerant and non-tolerant mice^94^. Comparisons of high-level relationships revealed that the connections among the most central nodes that represent keystone species (community members that connect the most community members) were significantly weaker and at times eliminated by chronic morphine, but these changes were not driven by individual genera within each central node (Figure 2B, detailed in Supplemental Table 3). Subsequent examination of pair-wise community member associations from these networks further reflected how morphine weakened or caused a loss of many associations (Supplemental Table 3). One positive association between two potential pathobionts, *Desulfovibrionaceae* and *Bacteroidales,* became stronger and one association between related Gram-positive taxa, an unclassified member of family *Erysipelotrichaceae* and genus *Coprobacillus,* was gained.

Since morphine experience in the paradigm appears to be a strong driver of microbiome assemblages, we evaluated whether morphine-driven disturbances were predictive of morphine use thereby strengthening inferences from microbiota associations. We did this by determining how well combinations of associated taxa identified from our experimental cohort could distinguish between pre-morphine versus morphine-treated microbiota from a different cohort of mice that experienced the same self-administration paradigm (Figure 1A). To evaluate this, we trained models on 30,295 different combinations of up to three community members associated with mid-to late morphine with our experimental cohort of mice (bred at Jackson Laboratory, JAX). We then tested these models on microbiota from a second cohort of mice bred in-house (n=4 female, n=4 male) (Figure 2C, data and statistics summarized in Supplemental Table 4 & 5). Even though the starting composition of the two experimental cohorts differed substantially, likely reflecting breeding facility, predictive models using the top eight ranking community profiles accurately identified most pre-morphine microbiota (Figure 2C). Furthermore, the models also had similar predictive power on all morphine-exposed microbiota of the in-house-bred mice compared to JAX-bred mice (46% as opposed to 57%). The robustness of the models across independent experiments throughout the paradigm suggests that nuances in starting microbiota composition did not strongly impact global morphine-induced compositional changes, even those that were sustained after morphine was removed, and that morphine induced common, predictive changes in the microbiota. This also exemplifies how co-occurrence indicator analyses could prove useful for identifying morphine use.

### Morphine concurrently depleted mutualistic community members that support gut homeostasis and increased relative abundance of pathobionts

Using taxonomic assignments, we identified the genera that were altered by morphine and best explained differences between pre-morphine and morphine-exposed microbiota, representing biomarkers of morphine exposure^88^. We then visualized temporal changes in their abundance during the different phases of morphine experience (see Figure 1A) compared to pre-morphine to glean insight into the dynamics of morphine dysbiosis (Figure 3A, Supplemental Table 2). Among 33 taxa identified as biomarkers, most were depleted by morphine and only emerged as biomarkers after prolonged morphine exposure. Fewer taxa increased in abundance with morphine exposure (Figure 3A, Supplemental Table 2). All biomarker taxa were also differentially variable in response to morphine (e.g. Supplemental Figure 4).Congruent with the expectation of morphine-driven dysbiosis, some potential pathobionts, including *Allobaculum, Prevotella, Erysipelotrichaceae,* and *Streptococcus*^108–113^, increased in abundance with morphine treatment, whereas some potential mutualistic taxa, including *Akkermansia*, *Bacteroides, Clostridium,* and *Rosburia*^26, 114, 115^ decreased in abundance (Figure 3B, Supplemental Table 2). Notably, many of the microbiota depleted by morphine are associated with gut barrier integrity, and/or are predicted SCFA-producing taxa^26, 116–120^ (Figure 3B, Supplemental Tables 6 & 7). Though these potentially beneficial genera trended towards a gradual depletion, some increased in abundance at times, including *Clostridium, Parabacteroides, Lactobacillus* and *Blautia*^116, 121, 122^ (Figure 3A, Supplemental Table 6 & 7).

**Figure 3:**
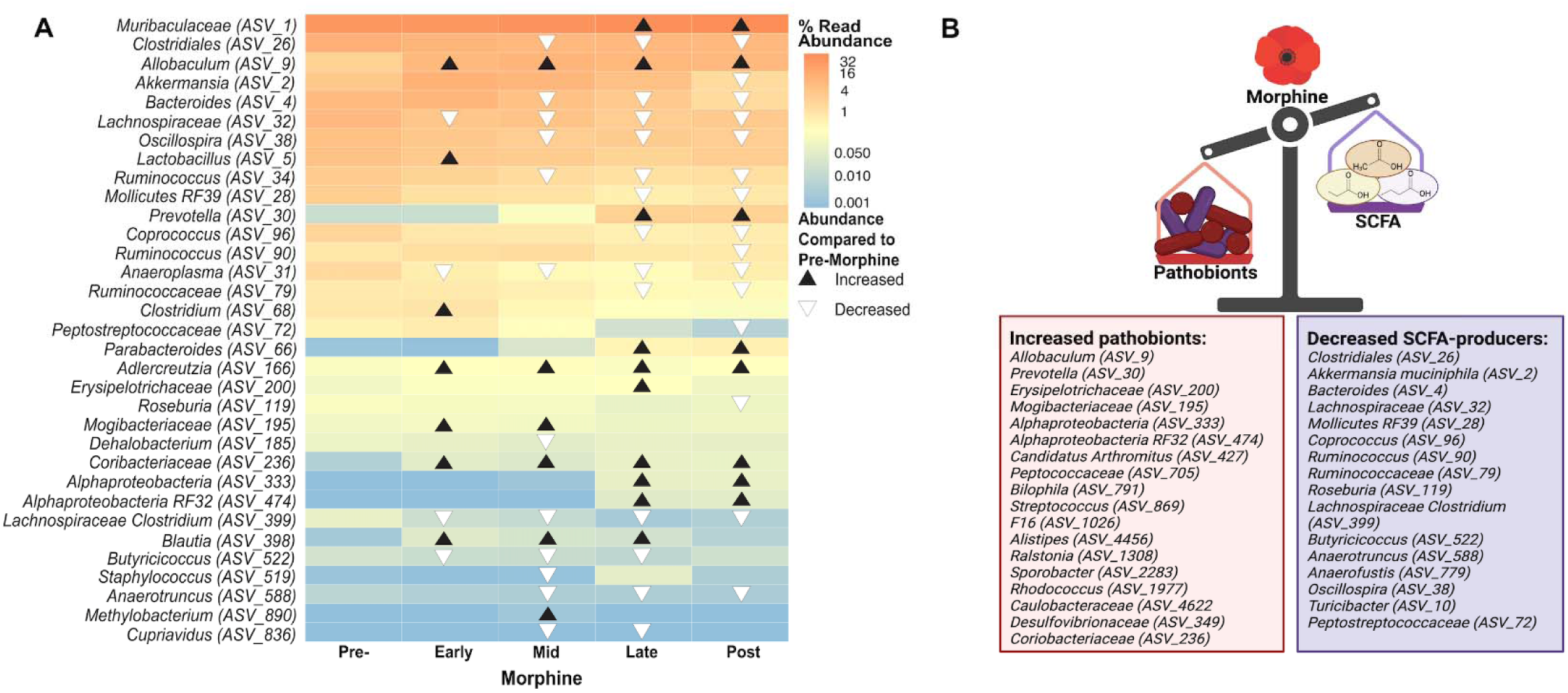
Morphine led to concurrent increase in pathobionts and decrease in mutualists that produce short chain fatty acids (SCFAs). A) Biomarkers of morphine consumption were identified from among the community members whose abundance was significantly altered by morphine (37 species, ∼21% of taxa) using corncob binomial regression models^88^, followed by a linear discrimination analysis (LDA) of effect size (LEfSe)^89^. Triangles indicate where in the paradigm the community member was a biomarker and whether its abundance increased (black) or decreased (white) compared to pre-morphine. Community members are labeled at the lowest taxonomic classification available. See Supplemental Table 2 for details. B) Functions of the microbiot that were differentially abundant during and after morphine exposure relative to their starting abundances were inferred from published studies (See Supplemental Table 7). Figure created with BioRender.com.

Collectively, these analyses highlight the significant, progressing, and enduring effects of prolonged morphine use, where the microbiota did not return to its pre-morphine exposure state, at least not within the several weeks after morphine was removed (see Figure 1A for paradigm details). These changes also align well with evidence that morphine-driven dysbiosis precipitates tolerance via pathobiont overgrowth which has the potential to trigger uncontrolled inflammation due to loss of commensals that help to attenuate this inflammatory response ^10, 37–42^. However, morphine produced significant, progressing dysbiosis in all mice, even those that did not develop tolerance and, as such, tolerance cannot be explained simply by these global changes. We posited that if nuanced differences in the responses of the microbiota contributed to differences in the development of tolerance, they would likely be overshadowed unless the analyses accounted for the common morphine experience through time.

### Tolerant mice exhibited earlier microbiota instability and depletion of beneficial taxa

Even with the dominating morphine-induced dysbiosis in all mice, there were clues that morphine disturbed the microbiota differently in tolerant and non-tolerant mice. We explored this potential using a Bayesian methodology to examine change-points in α-diversity. This analysis revealed a significant decrease in count and balance of ASVs, and this occurred earlier, by week five of morphine exposure in tolerant mice as opposed to week 9 in non-tolerant mice (Figure 4A). Analysis of β-diversity also revealed that the microbiota of tolerant and non-tolerant mice differed in membership and abundance during morphine consumption (*p*_PERMANOVA_ = 0.0022, weighted UniFrac), though diversity did not differ either before or after morphine (Supplemental Table 1). These results suggest a link between the progression of microbiome dysbiosis and the difference in tolerance. The dramatic drop in diversity identified in microbiota from tolerant mice were detected during the mid-morphine phase of exposure but may have resulted from an earlier loss of stabilizing community members, as evidenced by divergences in diversity during the early morphine phase of exposure (Figure 4A).

**Figure 4:**
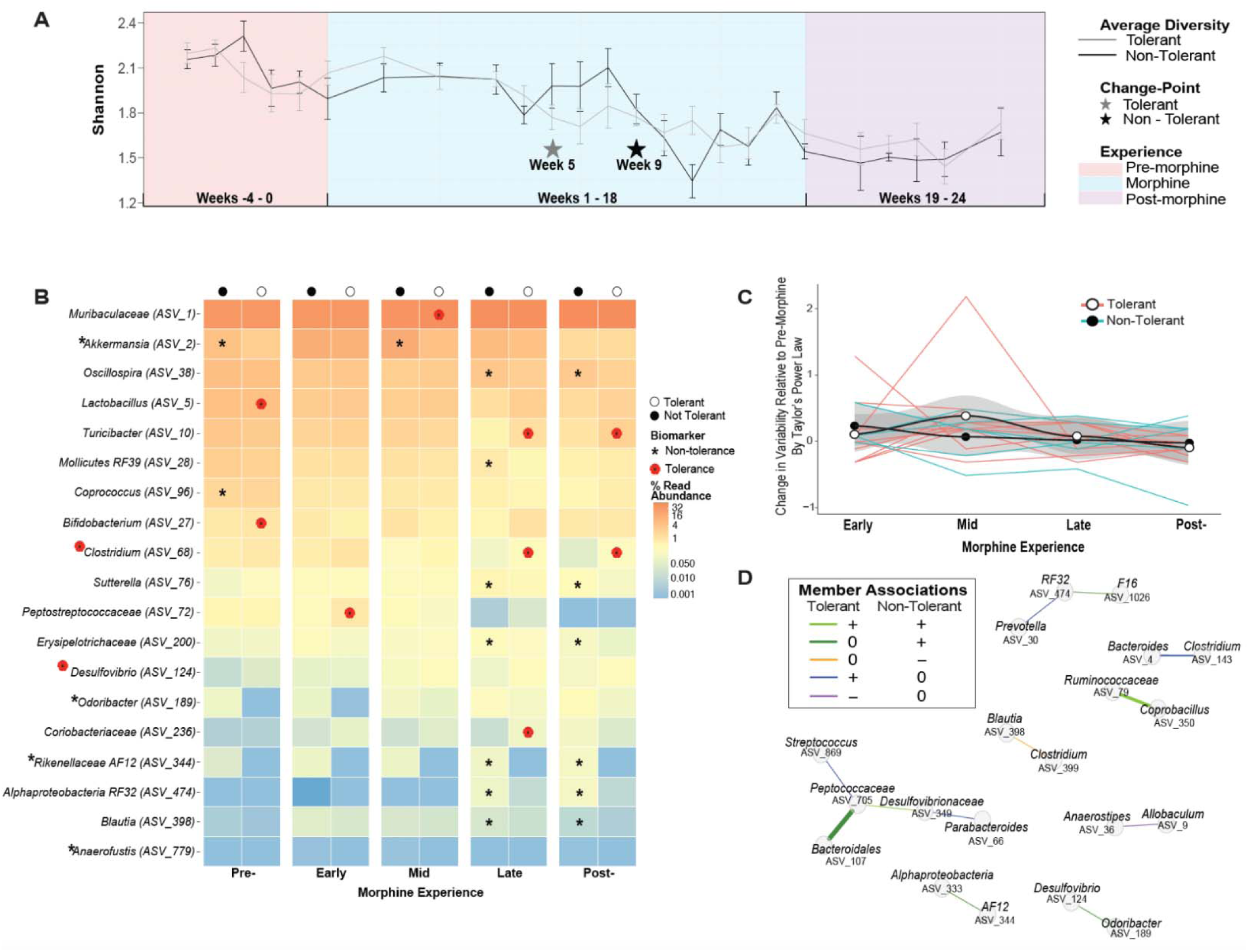
Analysis of differential responses of the microbiota to morphine disturbance reveal microbiota associations with variability of tolerance. A) Change in average microbiota α-alpha diversity at the genus level is plotted sequentially by sample number for tolerant (grey line) and non-tolerant (black line) mice, assessed using number and evenness of species (Shannon). Bayesian change-points in diversity of tolerant (grey star) and non-tolerant (black star) mice are shown. Error bars represent SEM. B) Biomarkers of antinociceptive tolerance to morphine were identified from among the community members whose abundance was explained by tolerance (as identified using corncob regression models)^88^, followed by a linear discrimination analysis (LDA) of effect size (LEfSe)^89^ representing genera that most likely explain differences between the microbiota of tolerant (poppy symbol) or non-tolerant (asterisk) mice. Community members are labeled at the genus level or at the lowest taxonomic classification available. *Desulfovibrio*, *Odoribacter*, and *Anaerofustis* were only identified as biomarkers when microbiota from all experiences were combined. See Supplemental Table 8 for details. C) Temporal stability of the gut microbiota of individual mice at each phase of morphine experience was assessed by analysis of Type IV Taylor’s Power Law parameters^91, 92^. The average slope parameter for each mouse at each phase of morphine exposure was normalized by subtracting the average pre-morphine slope parameter. Positively increasing average change in slope (black line with grey confidence intervals) represents microbiota instability. See methods for details and Supplemental Figure 7 for Type I analysis representing normal stochasticity. D) Differences in network associations between non-tolerant and tolerant mice. Associations of genera that were different between tolerant and non-tolerant mice, extracted from Supplemental Figure 8 are represented by a connecting line where colors indicate positive or negative associations and thickness indicates relative difference in strength of connection between non-tolerant and tolerant mice. See Supplemental Table 10 for statistics.

We next identified differentially abundant genera associated with the development of tolerance/protection from tolerance using a beta-binomial regression model to control for phase of morphine experience^88^. From among these genera, we identified biomarkers-genera that are significantly correlated with and most explained the difference between tolerant and non-tolerant mice^89^. These analyses revealed relatively few biomarkers, eight for tolerant mice and 11 for non-tolerant mice (Figure 4B, see Supplemental Table 8 for details and statistics). Three genera, all recognized as “probiotic,” were biomarkers before morphine exposure. Specifically, tolerant mice initially harbored higher abundances of *Lactobacillus* and *Bifidobacterium,* whereas non-tolerant mice exhibited higher abundances of *Akkermansia muciniphila* (Figure 4B). Although *A. muciniphila* increased after morphine exposure in tolerant and non-tolerant mice alike, a higher level was maintained longer in non-tolerant mice (Figure 4B, Supplemental Figure 5). Earlier changepoints, or periods of instability, led to some divergent patterns of abundance in tolerant versus non-tolerant mice, including higher abundance of pathobiotic genera *Desulfovibrio* and *Coriobacteracea*, and lower abundance of potentially beneficial taxa *Rickenellaceae* AF12 and *Odoribacter* in tolerant mice (Figure 4B, Supplemental Figure 5).

Most biomarkers were only apparent after morphine had substantially altered microbiota composition, with notable exceptions (e.g. *A. muciniphila*), but this does not preclude that individual microbiota differences earlier in the paradigm, or even prior to morphine exposure, impacted antinociception and/or the trajectory of morphine tolerance. For example, four mice exhibited low morphine antinociception prior to chronic morphine consumption and antinociception in these mice was further compromised by chronic morphine reflecting tolerance (Figure 1B, colored mice). The most noteworthy difference between the pre-morphine microbiota of mice with high and low starting antinociception was the near absence of *Odoribacter*^144^ in mice with low starting antinociception (Supplemental Figure 6; detailed statistics in Supplemental Table 9).

We also evaluated the stability of the gut microbiota of individual mice as a measure of dysbiosis and assessed whether the two populations-tolerant and non-tolerant mice-met the expectations of Taylor’s power law^91, 168, 169^. Taylor’s power law describes the scalable relationship between the variance in abundance of a taxon to its population mean and it can be used to explore ecologically meaningful differences in population dynamics that result from external drivers and interactions among different taxa^92, 170^. Application of the Type I power law extension^92^ showed an expected correlation of variance of individual genera with their population means regardless of phase of morphine experience and development of tolerance as demonstrated by significant aggregation and linear clustering of data (Supplemental Figure 7). However, application of the Type IV power law extension, which examines individual communities over time^92^, detected differences in temporal stability between microbiota from tolerant and non-tolerant mice (Figure 4C, see methods for details). Specifically, the slope parameter of microbiota from tolerant mice increased (positive value) during mid-morphine indicating greater variance relative to starting variance, a result that was not driven by any one individual mouse. In contrast, the slope parameter of microbiota from non-tolerant mice did not increase (0 or negative value). The increase in population variance in tolerant mice indicates temporal instability among these populations (Figure 4C). Importantly, this trend coincides with the change in α-diversity observed in tolerant mice at week 5 of the paradigm (Figure 4A).

Since morphine significantly altered relationships of the most central community members in all mice (Figure 2B), we wondered whether the relationships of microbiota of mice that did or did not develop tolerance differed^94^. Overall microbiota connectivity did not differ between tolerant and non-tolerant mice, further reinforcing that all mice experienced dysbiosis (Supplemental Figure 8). However, network differences between tolerant and non-tolerant mice were revealed by an assessment of whether the abundance of paired community members correlated with each other and were different in the two populations (Figure 4D, Supplemental Table 10). This analysis identified six stronger positive associations and two unique negative (inverse) correlations in mice that did not develop tolerance, and four positive associations and one negative correlation that were unique to tolerant mice.

We also interrogated whether patterns in unique membership, or indicator genera were linked to tolerance^90^. Only three taxa altered by morphine were differentially present in tolerant mice, and these were identified late and post-morphine or when all phases of morphine experience were combined (Table 1). In contrast, nine indicator taxa were identified from non-tolerant mice, some from every phase of morphine experience (Table 1). With the exceptions of *Odoribacter, Rickenellacea AF12, and Flexispira*, which are indicators of non-tolerance, and *Rhodococcus*, an indicator of tolerance, all indicator taxa were significantly impacted by morphine in terms of abundance (Figure 3A, Supplemental Table 2), as indicator species (Supplemental Table 4), or changes in network centrality (Figure 2C, Supplemental Table 3). *Bilophilia* was an indicator of both tolerance and morphine. *Streptococcus* was both an indicator of tolerance and displayed a unique network association in tolerant mice (Table 1, Supplemental Table 3, Supplemental Table 4).

**Table 1:** Indicator taxa of morphine tolerant and non-tolerant microbiota.

| Phase | Tolerant | Community Member | Specificity<sup>1</sup><br>Probability | Sensitivity<sup>2</sup><br>Probability | Indicator<br>Value<sup>3</sup> | p-value |
| --- | --- | --- | --- | --- | --- | --- |
| Pre | No | *Rikenellaceae AF12*<br>(ASV_344)<sup>4</sup> | 0.7955 | 0.1944 | 0.393 | 0.038 |
| Early | No | *Burkholderia* (ASV_1302)<sup>5</sup> | 0.8929 | 0.2778 | 0.498 | 0.024 |
|  | No | *Xanthomonas* (ASV_2124)<sup>5</sup> | 1 | 0.1667 | 0.408 | 0.045 |
| Mid | No | *Parabacteroides* (ASV_66)<sup>5</sup> | 0.7852 | 0.375 | 0.543 | 0.012 |
| Late | No | *RF32* (ASV_474)<sup>5</sup> | 0.7353 | 0.5556 | 0.639 | 0.001 |
|  | No | *AF12* (ASV_344)<sup>4</sup> | 0.8475 | 0.2778 | 0.485 | 0.004 |
|  | No | *F16* (ASV_1026)<sup>5</sup> | 0.7534 | 0.3056 | 0.48 | 0.018 |
|  | Yes | *Rhodococcus* (ASV_1977)<sup>5</sup> | 1 | 0.15 | 0.387 | 0.021 |
| Post | No | *Anaerofustis* (ASV_779)<sup>5</sup> | 0.7143 | 0.7143 | 0.714 | 0.001 |
|  | No | *Odoribacter* (ASV_189)<sup>4</sup> | 0.6829 | 0.5714 | 0.625 | 0.008 |
|  | No | *RF32* (ASV_474)<sup>5</sup> | 0.7538 | 0.5 | 0.614 | 0.005 |
|  | No | *F16* (ASV_1026)<sup>5</sup> | 0.7333 | 0.3929 | 0.537 | 0.015 |
|  | No | *Stenotrophomonas*<sup>5</sup><br>(ASV_7876) | 1 | 0.1071 | 0.327 | 0.047 |
|  | Yes | *Streptococcus* (ASV_869)<sup>5</sup> | 1 | 0.2245 | 0.474 | 0.02 |
| All | No | *RF32* (ASV_474)<sup>5</sup> | 0.70664 | 0.31081 | 0.469 | 0.001 |
|  | No | *AF12* (ASV_344)<sup>4</sup> | 0.73464 | 0.25676 | 0.434 | 0.001 |
|  | No | *F16* (ASV_1026)<sup>5</sup> | 0.71212 | 0.20946 | 0.386 | 0.002 |
|  | No | *Flexispira* (ASV_704)<sup>4</sup> | 1 | 0.02027 | 0.142 | 0.038 |
|  | Yes | *Bilophila* (ASV_791)<sup>5,6</sup> | 0.7886 | 0.1008 | 0.282 | 0.016 |

<sup>1</sup> Specificity is the estimate of the probability that the ASV belongs to the experience given the fact that the ASV was found. This conditional probability is called the *specificity* or *positive predictive value* of the species as indicator of the site group
<sup>2</sup> Sensitivity is sample estimate of the probability of finding the ASV in samples belonging to the experience. This second conditional probability is called the *fidelity* or *sensitivity* of the ASV as indicator of the target site group
<sup>3</sup> Indicator value index (accounts for unequal group sizes) given probabilities A and B
<sup>4</sup> Identified as a biomarker of non-tolerance but not influenced by morphine use (See Figure 4)
<sup>5</sup> Identified as biomarker, indicator species, or by centrality measures and linked to morphine use (See Figure 3A and Supplemental Table 3)
<sup>6</sup> Identified in higher abundance in mice with poor starting antinociception (see Supplemental Figure 5)

### Genera differentially associated with and predictive of non-tolerant mice were more stably maintained during morphine disturbance than genera associated with tolerant mice

To further examine the strength and patterns of microbiota associations with differences in the development of tolerance, we applied predictive models of abundance and presence of up to 3 community members identified at each phase of morphine experience in association with either tolerant or non-tolerant mice^90^(Summarized in Figure 5 and Supplemental Table 11, see Supplemental Table 12 for a full description models and statistics). Starting composition was not predictive of either outcome, in agreement with individual variability of pre-morphine microbiota (Figure 1D, Supplemental Figure 3), similar alpha and beta diversity, (e.g. Figure 4A) and the limited number of biomarkers identified pre-morphine (Figure 4B). The models built from morphine as well as post-morphine associations were more specific for non-tolerant microbiota and assigned these with an average of 86% accuracy, as opposed to only 45% accuracy for tolerant mice indicating distinguishing differences had developed in the microbiota of non-tolerant mice even during early morphine. By the post-morphine phase, models assigned microbiota from individual fecal samples to the tolerant and non-tolerant groups with 97% accuracy reflecting that the microbiota were divergent.

**Figure 5:**
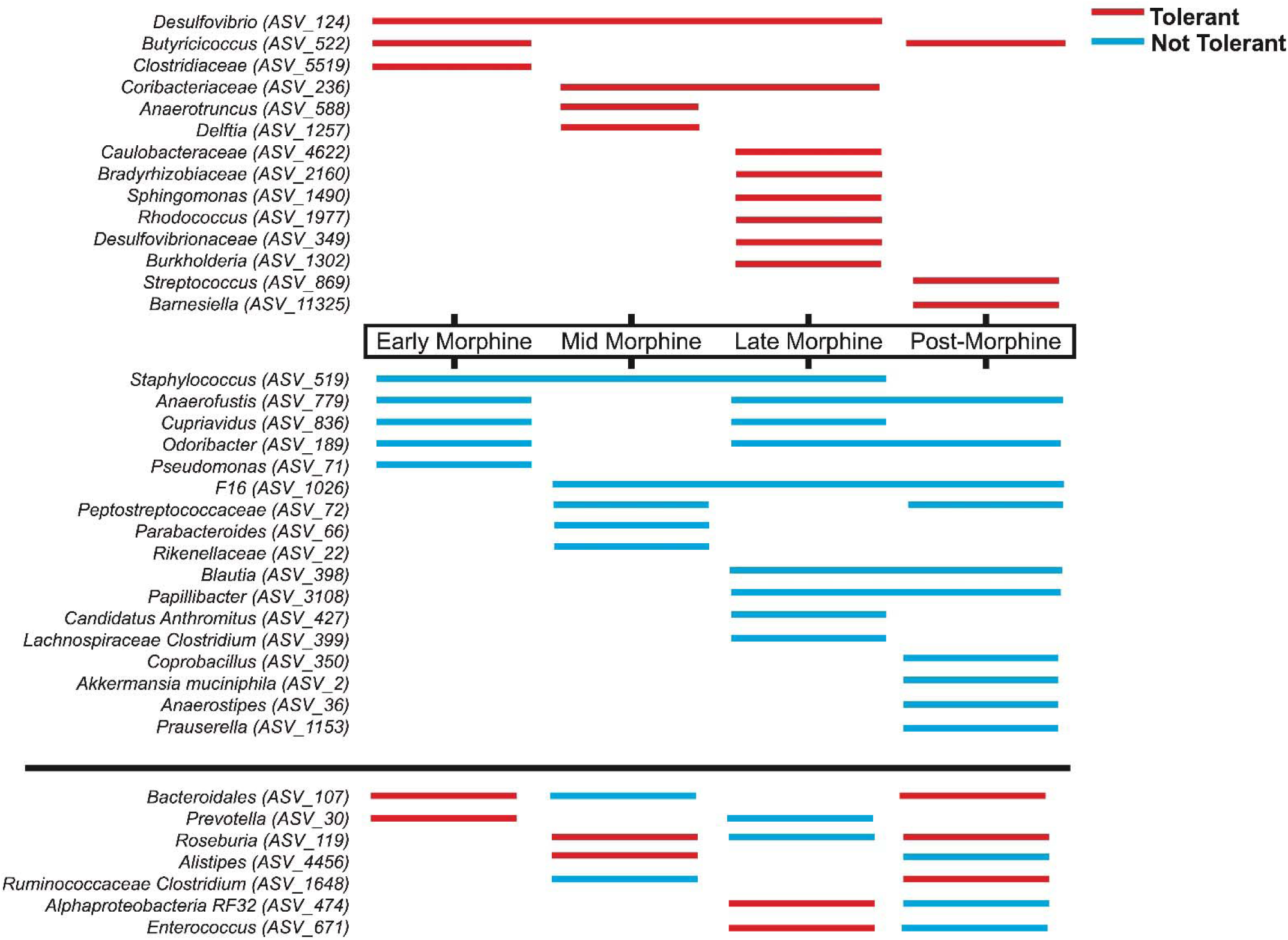
Summary of genera informing predictive models exemplifying more stable associations and shared predictive genera among non-tolerant than tolerant mice. Summary of taxa used in combinations to predict whether a microbiota sample was from a tolerant mouse (red) or a non-tolerant mouse (blue) in different phases of morphine experience. No models were predictive during pre-morphine. Three genera that were inconsistently identified in biomarker, indicator, or network analyses (showing the opposite associations in prior analyses at the same phase of morphine experience compared to these models) were excluded from this visual summary. See Supplemental Table 11 and 12 for details.

A visual exploration of the temporal distribution of the genera that informed these predictive models reveals that many genera predictive of non-tolerant microbiota spanned multiple phases of morphine experience, reflecting their stable association (Figure 5). In keeping with this, the statistically supported (p<0.05) non-tolerant microbiota models, built at discrete phases of morphine exposure, had more cross predictability of samples from other phases of the paradigm (68% of non-tolerant samples correctly identified versus 57% tolerant samples). Collectively, this data supports that microbiota from tolerant mice were more temporally unstable and more divergent from each other than the microbiota of non-tolerant mice. This is perhaps not surprising considering non-tolerant mice did not vary in antinociception, whereas tolerant mice varied more from one another (Figure 1B). But the cross-predictability between the microbiota of just 6 non-tolerant mice suggests a convergence in mechanisms of sustained antinociception as well as resistance to some morphine-driven changes (Figure 4).

Considering tolerant mice appear less alike each other and non-tolerant mice appear more convergent in shared and predictive genera, we re-evaluated similarity among microbiota that were from tolerant or non-tolerant mice and differences between these two populations using only biomarker genera and a phylogenetically informed metric of β-diversity (weighted UniFrac; Supplemental Figure 9). This analysis indicated that phase of morphine experience was still the major driver of assemblages, even when considering a restricted dataset (19 ASVs at the genus level). In addition, this analysis revealed more variability among the microbiomes of tolerant mice than non-tolerant mice during mid-morphine (Supplemental Figure 9). This agrees with the timing in which we predicted community instability in tolerant mice (Figure 4C) following their notable earlier changepoint in α-diversity (Figure 4A). In contrast, microbiota of non-tolerant mice clustered more tightly mid-morphine, suggesting more similarity and perhaps shared functions (Supplemental Figure 9).

### Greater relative microbiota capacity for butyrate production during morphine exposure correlated with prolonged morphine antinociception

The finding that mice that did not develop tolerance to morphine share more predictive genera and that associations were still quite different from each other in terms of community structure even when only considering biomarker genera (Figure 5, Supplemental Figure 9, Supplemental Table 8, and Supplemental Table 12), led us to consider whether functions among shared taxa could elucidate what these mice have in common. Assessment of previously described attributes of biomarker and indicator genera (Table 2) revealed a striking pattern among genera associated with non-tolerant mice: a higher abundance and association of butyrate producers that was corroborated by a bioinformatic analysis that predicts the functional capacity of taxa (Supplemental Figure 10). Congruent with this observation, butyrate bolsters gut barrier function and curtails inflammatory responses while stimulating macrophage differentiation and production of antimicrobial peptides to remove invading pathobionts, thereby promoting gut homeostasis^48, 171, 172^. It is notable that many of the butyrate producing genera associated with non-tolerant mice were also lower in mice with poor starting antinociception prior to morphine exposure and some were subsequently depleted even more by morphine, in parallel with the further decline of morphine antinociception in these mice (Figure 1B, Supplemental Figure 6). This suggests that morphine-driven differential loss of some butyrate-producing organisms might exacerbate tolerance and, by extension, that butyrate could protect against tolerance.

**Table 2:** Butyrate producers were associate with non-tolerant mice^1^.

| Taxa | Association<sup>2</sup> | Experience | Notable Feature(s) |
| --- | --- | --- | --- |
| *Akkermansia muciniphila* (ASV_2)<sup>3</sup> | B<br>A<br>M | Pre, Mid<br>Early, Late<br>Post | Mucin degrading, acetate producer; syntrophic support of **butyrate** producers<sup>117, 123</sup><br>Modulates immune responses and gut homeostasis<sup>124-128</sup> |
| *Oscillospira* (*Oscillibacter ruminatum*) (ASV_38)<sup>3</sup> | B<br>A | Late, Post<br>Early-Late/Post | **Butyrate**<sup>129</sup><br>Chronic constipation correlated with CD post-operative remission<sup>129, 130</sup> |
| *Rikenellaceae* (*Alistipes onderdonkii*) (ASV_22) | M | Mid | Potential acetate and propionate producer based on taxa<sup>131</sup><br>May affect tryptophan bioavailability in MDD<sup>132</sup> |
| Mollicutes RF39 (*Breznakia pachnodae*)(ASV_28)<sup>3</sup> | B | Late | Acetate producer; inversely correlated with inflammation, and BMI<sup>133, 134</sup> |
| *Coprococcus* (*Eubacterium xylanophilum*) (ASV_96)<sup>3</sup> | B | Pre- | **Butyrate** and acetate based on taxa; inversely correlates with IBD, depression and PD<sup>116, 135, 136</sup> |
| *Sutterella* (*Parasutterella excrementihominis*) (ASV_76) | B | Late, Post | Bile acid maintenance<sup>137</sup>; Morphine dysbiosis, inflammation<sup>138</sup>; Loperamide reduces butyrate production and *Sutterella*<sup>139</sup> |
| *Peptostreptococcaceae* (*Romboutsia timonensis*) (ASV_72) | M | Mid, Post | Mucin degrading, tryptophan metabolism and indole acrylic acid production promotes barrier function; controversially (positively/negatively) correlated with anxiety and depressive disorders<sup>140-143</sup> |
| *Parabacteroides* (*Parabacteroides chongii*) (ASV_66) | I<br>M | Early-Mid<br>Mid, All | **Butyrate**, anti-inflammatory<sup>121</sup> |
| *Odoribacter* (*Odoribacter splanchnicus*) (ASV_189) | B<br>A<br>I | Combined<br>Pre-Late/Post<br>Early, Late, Post | **Butyrate**, inversely correlated with gut inflammation, associated with exercise and weight control<sup>120, 144-147</sup> |
| *Erysipelotrichaceae* (*Clostridium*; *Longibaculum muris*) (ASV_143)<sup>3</sup> | B | Late, Post | Glucose absorption/glycemic control; Linked to intestinal inflammation<sup>111</sup><br>Potential butyrate producer based on taxa<sup>116</sup> |
| *Rikenellaceae* AF12 (*Millionella massiliensis*) (ASV_344) | B<br>A<br>I | Late, Post<br>Pre-Late/Post<br>Pre, Early-Mid | **Butyrate** producer associated with exercise<sup>144</sup> |
| *Alphaproteobacteria* RF32 (*Kiloniella laminariae*) | B<br>I | Late, Post<br>Early-Mid, Post | Colonic damage<sup>148</sup>, inverse with 5-HT<sup>149</sup> |

| (ASV_474)<sup>3</sup> | M<sup>4</sup> | Post |  |
| --- | --- | --- | --- |
| *Lachnospiraceae* (*Clostridium colinum*) (ASV_399) | M | Late, All | Potential **butyrate**, and Indoleacrylic acid producer based on taxa <sup>116, 140, 150</sup> |
| *Coprobacillus* (*Longibaculum muris*) (ASV_350) | M | Late | Positively correlated with leptin (anti-obesity) <sup>151</sup> |
| *Blautia* (*Murimonas intestini*) (ASV_398)<sup>3</sup> | B<br>A<br>I | Late, Post<br>Early-Post<br>Post | Acetate producer (Potential co-metabolism feeding for **butyrate** producers), inversely correlated with PD <sup>122</sup><br>Linked to gut homeostasis <sup>136, 152</sup> |
| *Staphylococcus* (*Staphylococcus saprophyticus*) (ASV_519) | A | Late | Grooming <sup>153</sup> |
| *Candidatus Anthromitus* (*Clostridium oryzae*) (ASV_427) | M | Late, Post | Potential **butyrate** producer based on taxa <sup>115, 116</sup><br>Linked to inflammation <sup>154</sup><br>Controversial taxonomic assignment <sup>155, 156</sup> |
| *Anaerofustis* (*Anaerofustis stercorihominis*) (ASV_779) | B<br>I<br>M | Combined<br>Post<br>Early, Late, Post, All | Acetate and **butyrate** <sup>157</sup> |
| *TM7 F16* (*Geomonas terrae*) (ASV_1026)<sup>3</sup> | I | Early-Mid, Post | Associated with gut dysbiosis and inflammation <sup>158, 159</sup> |
| *Cupriavidus* (*Cupriavidus metallidurans*) (ASV_836) | M | Early, Late | Environmental contaminant <sup>160</sup> |
| *Pseudomonas* (*Pseudomonas aeruginosa*) (ASV_71) | M | Early | Becomes more virulent in the presence of morphine <sup>161</sup> |
| *Papillibacter* (*Clostridium viride*)(ASV_3108) | M | Late, Post | Potential **butyrate** producer based on taxa <sup>115, 116</sup> |
| *Burkholderia* (*Burkholderia territorii*) (ASV_1302)<sup>3</sup> | I | Early | Potential pathogen <sup>162, 163</sup> |
| *Xanthomonas* (*Xanthomonas albilineans*) (ASV_2124)<sup>3</sup> | I | Early-Mid | Participate in bile acid metabolism <sup>164, 165</sup> |
| *Stenotrophomonas* (*Xanthomonas floridensis*)*(ASV_7876)<sup>3</sup> | I | Post | Associated with gut inflammation <sup>166, 167</sup> |

<sup>1</sup> Ordered by decreasing abundance. Species (in parentheses) were identified using rRNA/ITS databases with the nucleotide basic local alignment search tool (blastn) <sup>70</sup>.
<sup>2</sup> B: Biomarker of non-tolerance; A: High starting antinociception; I: Indicator taxa of non-tolerance; M: Predictive modelling.
<sup>3</sup> Altered by morphine exposure
<sup>4</sup> Identified in predictive models for both tolerance and non-tolerant mice but in combination with different taxa

To further investigate associations of the microbiota and its capacity to produce butyrate with sustained antinociception to morphine, we optimized a shorter paradigm for hypothesis testing that uses moderate but consistent non-contingent morphine exposure to produce tolerance (Figure 6A). This shorter paradigm more closely resembles a post-operative regime and eliminates the potentially confounding variable of different amounts of voluntary oral morphine on microbiome composition. While it accelerates the timeline for development of tolerance, it preserves variation in both microbiota composition and the development of tolerance (Figure 6A-B). Five days of morphine produced antinociceptive tolerance in most mice, with ∼30% of mice maintaining 100% MPE morphine antinociception (Figure 1B, 6B). The microbiota derived from feces pooled from non-tolerant and pooled from tolerant mice from this paradigm differed from each other post-morphine, suggesting microbiota differences could relate to variability in antinociceptive tolerance (Figure 6B).

**Figure 6:**
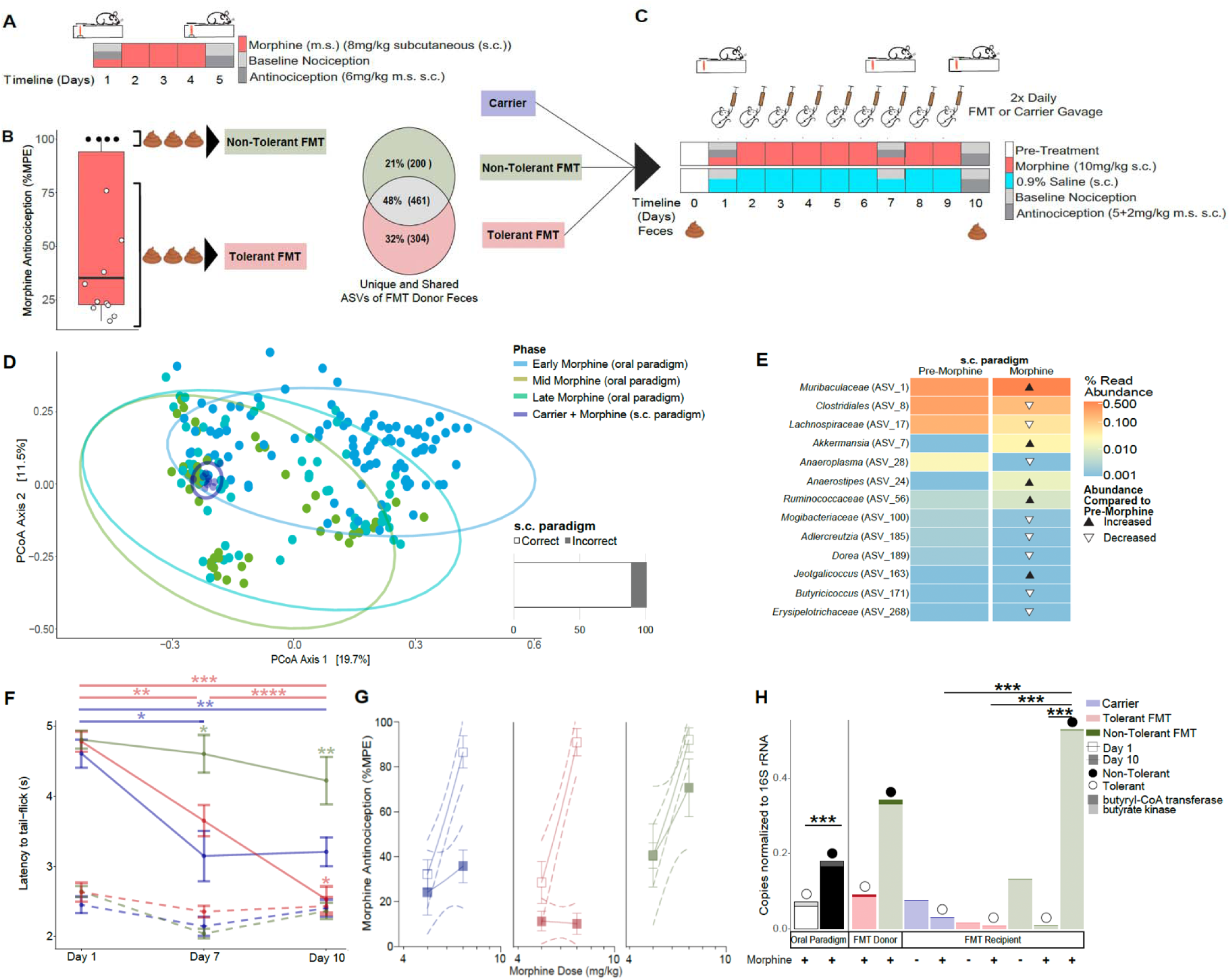
Microbiota enriched for butyrate biosynthetic capacity protected fecal microbiota transplantation (FMT) recipients from developing antinociceptive tolerance to morphine. A) Schematic of sub-cutaneous (s.c.) morphine (m.s.) (8 mg/kg m.s. s.c., 1x daily) paradigm. B) Baseline nociception and antinociception (6 mg/kg m.s.) was assessed on days 1 and 5 using a radiant heat tail-flick assay to identify tolerant (<100% maximum possible effect [MPE], light red) and non-tolerant mice (= 100% MPE, light green). Day-5 feces were pooled from each category for FMT, and for analysis of microbiota (16S) where compositional comparisons were illustrated using the ps_venn function in the MicEco R package (v0.9.15)^58^.C) Schematic of FMT paradigm where mice were orally gavaged 2x daily with saline carrier (dark blue), or cleared fecal slurries (See Figure 6B) in conjunction with daily s.c. m.s. administration (n = 9 ea treatment) or saline ( n = 5 ea treatment). D) Gut microbiota β-diversity (All ASVs, Bray-Curtis dissimilarity) of saline gavaged mice post-m.s. in comparison with microbiota from the oral paradigm as visualized by principal coordinates analysis (PCoA). The bar graph to the right shows accuracy by which the predictive models built from microbiota associations with m.s exposure in the oral paradigm (Figure 2C) could predict microbiota from the s.c. paradigm were derived from m.s treatment (presented as % correct/incorrect). See Supplemental Tables 5 and 13 for details. E) Biomarkers of s.c. m.s. exposure were identified from among the community members whose abundance was significantly altered by m.s. (as identified using corncob regression models)^88^, followed by a linear discrimination analysis (LDA) of effect size (LEfSe)^89^ of day 0 and day 10 microbiota. Triangles indicate whether the community members’ abundance increased (black) or decreased (white) day 10 relative to day 0. Community members are labeled at the lowest taxonomic classification available. See Supplemental Table 14 for details. F) The trajectory of tolerance was visualized as a decrease in latency to tail-flick in seconds following 5 + 2 mg/kg s.c. m.s. (cumulative ED_90_) and reported as mean; error bars are SEM. Kruskal-Wallis test followed by pairwise Wilcox test for multiple comparisons were used to determine whether carrier and FMT groups (solid lines) changed over time (horizontal bars) and differed from each other (asterisks above data points) (*p<0.05; **p<0.01; ***p<0.001; ****p<0.0001). Baseline latencies to tail-flick for carrier and FMT groups (dashed lines) were used to assess hyperalgesia post-chronic m.s.. G) Change in morphine potency was assessed by comparison of linear regressions of dose responses (ED_60_ and ED_90_ doses) reported as % MPE on day 1 (open squares) and day 10 (closed squares). Outer dashed lines represent 95% confidence intervals of the linear regressions and error bars represent SEM. Shift in ED_50_ from day 1 to day 10 extrapolated from these regressions is reported in text. H) Butyrate biosynthesis capacity determined by quantitative real-time polymerase chain reaction (qPCR) via gene copy number quantification of butyryl-CoA transferase (bcoat; upper stacked bar) and butyrate kinase (buk; lower stacked bar) normalized to total 16S rDNA copy number. Relative butyrate production capacity of tolerant (white bar) and non-tolerant (black bar) mice in the oral paradigm is presented in the first bar graph, followed by capacity in FMT donor slurries and recipients at day 10 for control carrier (blue), tolerant FMT (light red), and non-tolerant FMT (light green). Circles above the bars designate whether the mice that were tolerant (white) or non-tolerant (black) representing data from Figure 1B and Supplemental Figure 11B. Student’s T-test determined significant differences of butyrate capacity, (ns p>0.05; *p<0.05; **p<0.01, ***p<0.001).

To more directly examine the connection between the microbiota and variability in morphine tolerance we used tolerant and non-tolerant mice (Figure 6B) as fecal microbial transplantation (FMT) donors to investigate whether the microbiota of tolerant mice predisposed or accelerated the development of tolerance in recipients^40, 48^ and/or whether the microbiota of mice that did not develop tolerance could delay or prevent the development of tolerance in recipients. For these FMT experiments, mice received s.c. morphine (10 mg/kg) for 9-days to reliably produce tolerance in controls (receiving oral carrier, Figure 6C). After 9-days of morphine, the resulting microbiota of control mice receiving carrier (Figure 6D, dark blue circles) clustered tightly and generally among microbiota from the mid-morphine phase of the original oral paradigm. Furthermore, the predictive models built from microbiota associations with morphine exposure in the oral paradigm and that were able to detect fecal microbiota exposed to morphine from a separate cohort of mice treated with the same oral paradigm (Figure 2C, Supplemental Table 5) also correctly assigned eight out of nine feces from these morphine treated carrier-gavaged mice, reinforcing that 9-days of s.c. morphine produces similar dysbiosis (Figure 6D, Supplemental Table 13). The short paradigm also caused similar changes among many of the same community members (Figure 3A) (e.g. *Muribaculaceae, Clostridiales, Lachnospriaceae, Akkermansia, Anaeroplasma,* and *Butyricicoccus)* and a depletion of beneficial community members characteristic of morphine dysbiosis, notably some of which are butyrate producers (e.g. *Butyricicoccus,* Figure 6E, detailed statistics in Supplemental Table 14). Oral carrier-gavaged mice readily developed tolerance by day seven of daily morphine treatment as evidenced by their reduced latency to tail flick (Figure 6F) and decreased morphine antinociception at 5+2mg/kg which was the ED_90_ dose in morphine-naïve mice (Supplemental Figure 11A, B). Mice that received 2x daily tolerant mouse FMT similarly developed tolerance by day seven and displayed further profound loss of morphine antinociception at day 10 (Figure 6F & G, Supplemental Figure 11B). Remarkably, mice receiving 2x daily non-tolerant donor FMT during morphine treatment did not develop tolerance by day seven, and as a population their morphine antinociception did not differ on day 10 compared to day one, where five of the nine mice retained 100% MPE (Supplemental Figure 11B). Compared to control carrier mice that exhibited a significant decrease in morphine potency from day one (ED_50_ 5.66mg/kg) to day 10 (ED_50_ 9.47mg/kg, p=0.012), non-tolerant FMT mice exhibited no significant change in morphine potency from day 1 (ED_50_ 5.37mg/kg) to day 10 (ED_50_ 5.63mg/kg, p=0.34) (Figure 6G). Morphine-treatment mice had no increase in baseline nociception in any gavage group indicating they did not develop hyperalgesia (Figure 6H). Furthermore, 9-days of any FMT treatment without chronic morphine did not significantly alter baseline nociception or morphine antinociception (Supplemental Figure 11C, D).

Because the above data reinforces the link between differential morphine-induced changes in the microbiome with variability in the development of antinociceptive tolerance, and implicated butyrate producers, we used this paradigm to assess whether sustained antinociception was associated with butyrate production capacity. To this end, we quantified the relative abundance of genes encoding representative enzymes including butyryl-CoA transferase (BCoAT, *bcoat* or *but*) and butyrate kinase (*buk*), necessary for each of the two known pathways for butyrate biosynthesis^105, 106^, as a proxy of butyrate production capacity in the microbiota of tolerant and non-tolerant mice. We performed this analysis of microbiota of the individual mice from the voluntary oral paradigm (Figure 1A, B), of feces pooled from tolerant and pooled from non-tolerant mice used the as FMT donors (Figure 6A, B), and from the individual FMT/saline carrier recipients at day 10 (Figure 6F-G). This revealed that the microbiota of mice that did not develop morphine antinociceptive tolerance consistently had significantly higher capacity for butyrate production (Figure 6H). The ending microbiota (18 weeks) of non-tolerant mice from the oral paradigm had ∼26% higher butyrate production capacity than tolerant mice (Fig. 6H, oral paradigm). Pooled feces from non-tolerant mice from the s.c. paradigm used as FMT donors also had substantially higher butyrate production capacity than the pooled feces of tolerant counterpart mice (3.7-fold greater, Fig. 6H, “donor”). The non-tolerant FMT recipients that maintained 100% MPE antinociception had significantly higher butyrate production capacity than those that did become tolerant, including those that received non-tolerant FMT (23-fold greater), those that received a tolerant FMT gavage (60-fold greater), or a saline gavage (17.4-fold greater, Fig. 6H, “recipient”).

### Dietetic butyrate supplementation prevented the development of tolerance which arose without systemic inflammation or hyperalgesia

Since the short paradigm recapitulated the original microbiome associations and reinforces the hypothesis that butyrate is protective, we used this paradigm to evaluate the impact of dietetic butyrate supplementation on tolerance. Mice were supplied with sodium butyrate, or as a control sodium/pH-matched sodium citrate *ad libitum* in their drinking water for three weeks prior to morphine exposure, where we estimate mice consumed ∼3 g/kg butyrate daily. Day one latencies to tail flick without and with morphine did not differ between butyrate and citrate supplemented mice (Supplemental Figure 12A). However, only one butyrate mouse (∼6%) was below 100% MPE at the 5 + 2 cumulative morphine dose as opposed to four citrate mice (∼22%). This implies butyrate pretreatment may modestly bolster morphine antinociception.

Mice were then treated with non-contingent morphine (10 mg/kg s.c. 1x day for 9 days) or saline while continuing to receive supplementation (Figure 7A). On day 10, the baseline nociception of neither supplementation group treated with morphine differed from their respective saline control groups and few individuals exhibited an increase in nociception which also did not correlate with decreased antinociception indicating these mice did not develop hyperalgesia (Supplemental Figure 12B,C). As expected, morphine modestly reduced gut transit, as assessed by an increase in production of fecal pellets by reversal with the MOR antagonist naloxone, in all morphine-treated mice, indicating that butyrate did not appear to counter morphine-driven constipation (i.e. reduced gut motility) (Supplemental Figure 12D) as previously shown ^48^.

**Figure 7:**
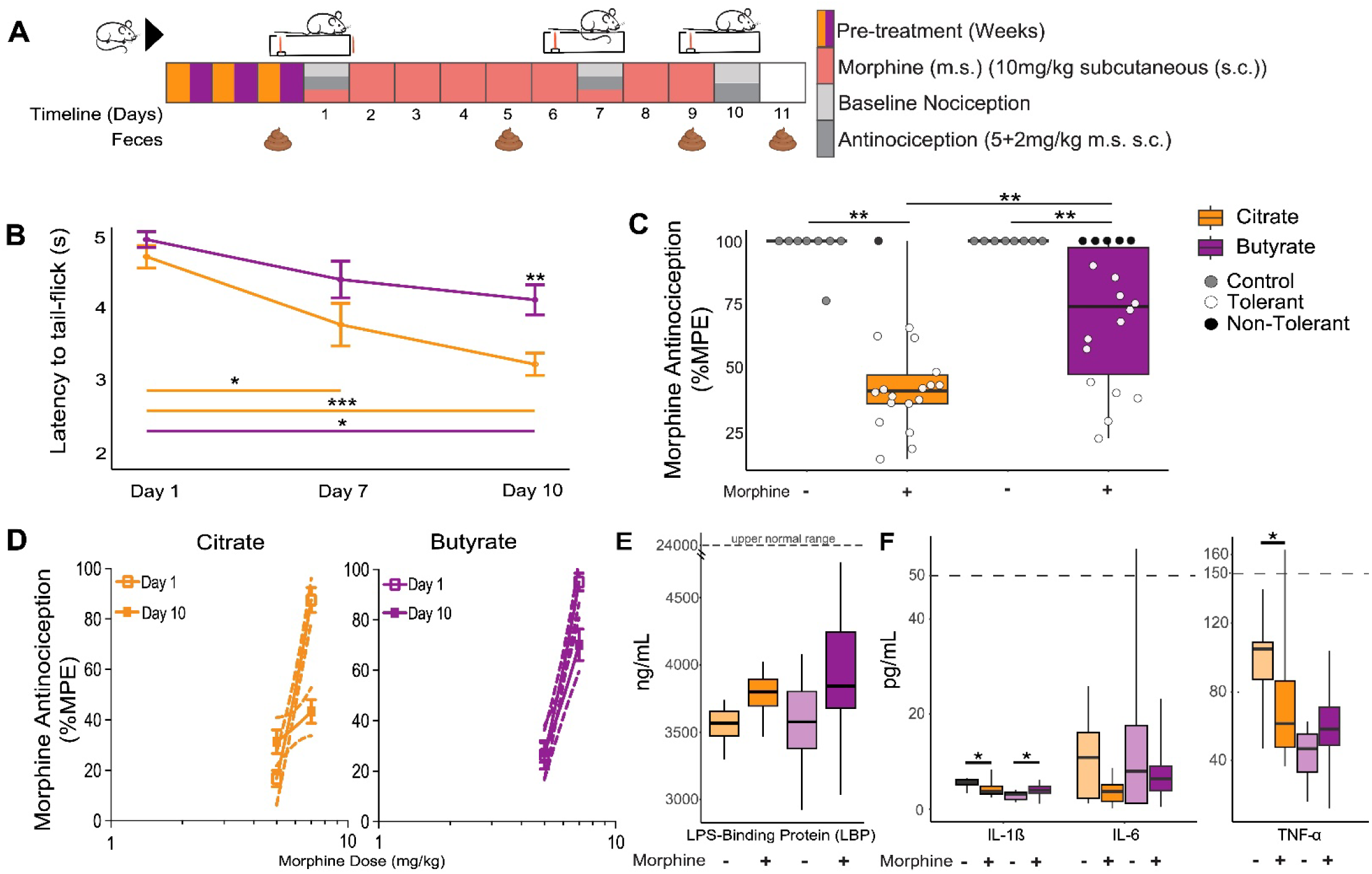
Butyrate supplementation prolongs morphine antinociception independently of curtailment of inflammation. A) Schematic of mouse paradigm where mice were fed 100 mM sodium butyrate or sodium citrate, followed either by 9-days morphine (m.s.) (10 mg/kg s.c.) (n = 18 ea), or daily saline (0.9% s.c.) (n = 8 ea). Mice treated daily with m.s. were assessed for baseline nociception and antinociception on days 1, 7, and 10 at 5mg/kg and 30 minutes later at 5+2 mg/kg m.s. s.c. (ED_60_ and ED_90_ respectively, see Supplemental Figure 11A). Feces were collected on Days 1, 5, 9, and 11 of the paradigm to generate 16S (V4-V5 regions, see methods). B). The trajectory of tolerance was visualized as a decrease in latency to tail-flick (s) (5 + 2 mg/kg s.c. m.s.) and reported as mean; error bars are SEM. Kruskal-Wallis test followed by pairwise Wilcox test for multiple comparisons were used to determine whether citrate and butyrate groups changed over time (horizontal bars) and differed from each other (asterisks above data points) (*p<0.05; **p<0.01; ***p<0.001). C) Antinociception (5 + 2 mg/kg s.c. m.s.) as determined on day 10 using the radiant heat tail flick assay. Data is reported as maximum possible effect (% MPE). Kruskal-Wallis test followed by pairwise Wilcox test for multiple comparisons determined significant differences between m.s. treated citrate and butyrate supplementation groups and between m.s. treatments and saline controls (**p<0.01; ***p<0.001). D) Change in morphine potency was assessed by comparison of linear regressions of dose responses (ED_60_ and ED_90_ doses) reported as %MPE on Day 1 (open squares) and Day 10 (closed squares). Outer dashed lines represent 95% confidence intervals of the linear regressions and error bars represent SEM. Shift in ED_50_ from day 1 to day 10 extrapolated from these regressions is reported in text. E) Day 10 serum levels of LPS binding protein (LBP) of mice supplemented with citrate (orange) or butyrate (purple) and their saline controls. LBP upper normal range (dashed line), as indicated by manufacturer, is marked for reference (Invitrogen, Waltham, MA). F) Day 10 serum levels of pro-inflammatory cytokines (IL-1β, IL-6, and TNF-α) of mice supplemented with citrate (orange) or butyrate (purple) and their saline controls. Dashed lines represent normal ranges of serum cytokine levels (50 pg/mL: IL-1β, IL-6; 150 pg/mL: TNF-α)^175^. Kruskal-Wallis test followed by pairwise Wilcox test for multiple comparisons determined significant differences between m.s. treated citrate and butyrate supplementation groups and between m.s. treatments and saline controls (*p<0.05).

Morphine-mediated antinociception declined precipitously in citrate-supplemented mice, and as a population only citrate-supplemented mice were tolerant by day seven (Figure 7B). Furthermore, on day 10, butyrate-supplemented mice maintained significantly higher morphine antinociception than citrate-supplemented mice (Figure 7C). More specifically, only citrate-supplemented mice exhibited a significant decline in morphine potency from ED_50_ of 5.9mg/kg to 8.1 mg/kg (p=0.0001), whereas the butyrate-supplemented mice exhibited a significantly smaller decrease in morphine potency (from ED_50_ 5.46 mg/kg to 5.88mg/kg,p=0.0028). Day 10 citrate vs. butyrate) (Figure 7D). Furthermore, both citrate and butyrate similarly altered the microbiome composition of mice (Supplemental Figure 12E), and as expected^48^, s.c. morphine induced dramatic microbiota changes even with butyrate supplementation (Supplemental Figure 12F, statistics detailed in Supplemental Table 15). Even so, morphine drove less dramatic changes in microbiota diversity and caused less instability in the microbiota of butyrate-than in citrate-supplemented mice (Supplemental Figure 12G, H).

Finally, we evaluated whether butyrate supplementation during morphine exposure improved barrier function by assessing serum LPS and LPS-binding protein (LBP) produced by the liver in response to bacterial translocation as surrogates of a “leaky gut.” We also assessed whether butyrate during morphine exposure curtailed systemic inflammation through quantification of blood serum inflammatory cytokines. In the short s.c. tolerance paradigm, the modest morphine dosing did not drive detectable increases in serum LPS, which were at or below the detection threshold (data not shown) indicating the mice were not septic^173^. Morphine also did not significantly elevate LBP levels, which were well below those expected if bacterial translocation had increased^174^ and these levels did not differ between supplementation groups (Figure 7E). In keeping with the low observed bacterial translocation, neither citrate-nor butyrate-supplemented mice displayed a notable increase in pro-inflammatory cytokines (IL-1β, IL-6, TNF-α) in serum post-morphine, and levels were all within the expected normal range (Figure 7F, dashed line). Indeed, the level of these were at times lower post-morphine compared to control mice that did not receive morphine perhaps because of the anti-inflammatory properties of morphine. This data indicates that in this paradigm, systemic inflammation was not required for the development of tolerance. It also suggests that paradigms that use super-physiological morphine doses may create phenomena that are less relevant to the clinical phenomenon.

## DISCUSSION

Prior studies examining contributions of the microbiome to OUD have intentionally reduced behavioral variation inherent to mouse models by using high non-contingent morphine doses. These prior studies have thereby gleaned insight from common microbiome responses, not all of which are likely to explain tolerance^17, 37, 40^. In contrast, through our experimental design, we preserved and leveraged natural variation in both tolerance and the microbiota with the specific goal to identify microbiota signatures associated with degree of tolerance (Figure 1 and 6). We demonstrate that chronic voluntary morphine self-administration induced similar, even predictive, dysbiosis in all mice, regardless of whether they develop antinociceptive tolerance (Figure 3A, Table 1), and these global temporal changes largely masked informative microbiome associations that tracked with differences in morphine tolerance. To uncover latent patterns associated with tolerance, we used complimentary high resolution temporal analyses and cross-referenced results to identify repetitive trends. These analyses revealed a divergence in the timing of progression of dysbiosis between tolerant and non-tolerant mice (Figure 4). More importantly, there were noteworthy differences in the response of microbiota to morphine disturbance that best distinguished between mice that did and did not develop tolerance (Figure 4, Figure 5, Table 2). Key among these findings was that non-tolerant mice maintained higher capacity for production of butyrate, (Table 2, Figure 6H)^8, 175^. We then showed that both FMT from morphine-treated non-tolerant donor mice relatively enriched for butyrate production capacity, and diet supplementation with butyrate protected against the development of tolerance to morphine (Figure 6F-H, & 7). Although butyrate is known to bolster barrier function and suppress inflammation, our data indicate that our moderate morphine tolerance paradigm does not produce systemic inflammation nor compromise barrier function (Fig. 7E and F). This uncoupling of tolerance from systemic inflammation and the ability of butyrate, either produced from the microbiota or as a dietary supplement, to reduce tolerance suggests a role of butyrate as a ligand of communication in the GMB axis independent from these palliative effects. (Figure 6 & 7).

As researchers increasingly recognize the importance of the gut microbiota and its signaling through the GMB-axis to host health and disease^1–4^, there has been a shift in thinking about how hosts and their associated microbial communities, known as the holobiont, interact to influence individual phenotypes^176–178^. Prior studies suggest that the gut microbiota enhances morphine tolerance through its translocation across a morphine-compromised gut barrier, as tolerance can be reduced by elimination of the gut microbiota via antibiotics, or in germ free mice^37, 40, 109^. The implication that the microbiota is the problem paints an incomplete picture, especially considering that depletion of gut mutualists, and their supportive functions, is arguably the most apparent feature of morphine-induced dysbiosis (Figure 3)^138^. Furthermore, a diverse gut microbiota is crucial for both immune and neurological functions^3, 179–181^. Indeed, supplementation with a probiotic mix of *Lactobacillus* and *Bifidobacteria*, gram positive genera shown to be depleted by morphine^38, 39, 41^, improves barrier integrity and attenuates morphine tolerance ^40^. Oddly enough, higher populations of these same genera were biomarkers of tolerance in our study here (Figure 4B) and *Lactobacillus* modestly increased in response to morphine in tolerant but not in non-tolerant mice (Figure 3A). Depletion of gram-positive bacteria with vancomycin, which targets these probiotic genera also prevents morphine tolerance^37^. We expect this incongruence, whereby these beneficial genera are tied to both tolerance and its attenuation, could reflect that a variety of taxa, not just these two genera, can support homeostasis similarly and support trophic networks of the microbiota to resist or counter morphine’s adverse effects (Figure 2D, Table 2). Importantly, these capabilities may not necessarily be defined by or limited to related taxa, or even guaranteed to be expressed equally by every strain within a given species (Supplemental Figure 6 and Table 2)^182, 183^. Functional redundancy of SCFA production by the microbiota could ensure resiliency to perturbation^184–187^.

Among SCFAs, which include acetate, propionate, and valerate, butyrate is especially important in the intestinal environment as the primary energy source for colonocytes, for promoting barrier function, and for toning the immune system to effectively mitigate translocating bacteria if barriers are breeched^48, 171, 172, 175, 188^. Notable potential butyrate producers associated with non-tolerance included *Oscillospira*, *Coprococcus*, *Parabacteriodes*, *Anaerofustis*, *Odoribacter, and Rickenellaceae AF12,* (Table 2)^116, 120, 121, 144–146, 150, 157, 189–193^. Many of these genera have been shown to improve gut integrity and inversely correlate with escalatory inflammatory conditions such as inflammatory bowel disease, cardio-metabolic diseases, and neurobiological or neurodegenerative conditions^120, 121, 125, 136, 144–146, 194–199^. Opioid use depletes butyrate-producing genera in humans^200^ and drives a decrease in fecal butyrate in mice ^48^. Curiously, the morphine agonist loperamide also decreases fecal butyrate even in the absence of constipation directly implicating MOR-signaling in microbiome dysbiosis ^139^. The replication of protection from tolerance by FMT enriched in butyrate biosynthesis capacity and diet supplementation of butyrate alone (Figure 6 & 7) suggests differential loss of such community members contributed to tolerance but did so without causing hyperalgesia and independently of cytokine-directed systemic inflammation that has previously been a suggested cause of microbiome-mediated hyperalgesia and tolerance^17, 37, 40^. Though the modest morphine regime we employed did not drive systemic inflammation or hyperalgesia (Supplemental Figure 2A, Figure 7, and Supplemental Figure 12B,C), we did not assess local intestinal inflammation which could still contribute to the pathology of tolerance. It is possible that butyrate concurrently palliatively counters the reduced analgesic benefits of morphine as tolerance develops by attenuating local inflammation, thereby preventing hyperactivation of nociceptor neurons that infiltrate the gut which could underlie hyperalgesia^40, 48, 201^, and which aligns with a recent study showing butyrate protection from hyperalgesia^46^. Importantly, the modest morphine regimes used herein uncoupled potential confounding side-effects that associate with tolerance when higher than necessary dosing of morphine is used, and these could be confounders resulting from experimental design that are neither required, nor the sole basis for microbiome-mediated exacerbation or protection.

As a complementary mechanism to its documented ability to reduce inflammation, butyrate could have modulatory effects on neuronal activity directly that may help explain its protective effects against the development of tolerance that developed in control mice independently of systemic inflammation (Figure 7). The gut is innervated by both enteric and sensory nerves, and both neuronal populations express opioid receptors^202^. When activated, opioid receptors decrease neuronal excitability and reduce neurotransmitter release. Neurons expressing opioid receptors in both the gut and the dorsal root ganglion (DRG) undergo homeostatic adaptations to chronic morphine that increase baseline neuronal activity^203, 204^. This neuronal “superactivation” in response to chronic opioid manifests as tolerance in the presence of drug, and withdrawal signs, including hyperalgesia, upon removal of opioid^30^. Butyrate could change the course/severity of the homeostatic adaptations in neurons thought to underlie tolerance and/or hyperalgesia associated with chronic morphine administration. In support of a direct role of butyrate on neuronal excitability, co-administration of butyrate in morphine-treated mice prevents morphine-induced changes in excitability of opioid receptor expressing neurons in the DRG^46^ thought to underlie hyperalgesia. In addition, as a ligand for two of its receptors, including the free-fatty acid receptor 3 (FFAR3) and GPR109A, which, like MOR, are Gi-coupled receptors, butyrate could mask or prevent hyperexcitability by providing Gi signaling. These are not the only possibilities. For example, the microbiota has also been shown to alter brain levels of brain-derived neurotrophic factor (BDNF) independent of the vagus nerve^205^, indicating the existence of microbial-produced signals that can cross the blood brain barrier beyond the DRG. SCFAs are one of several such microbial compounds known to be transmitted through the circulatory system that cross the blood-brain barrier ^6, 206, 207^. Because butyrate was protective for tolerance even in the absence of concurrent effects tied to systemic inflammation, butyrate supplementation could be a useful tool for experimentally identifying and uncoupling parallel mechanisms that contribute to antinociceptive tolerance^46^.

The most abundant biomarker of non-tolerance, *A. muciniphila*, was a particularly intriguing association due to its role in promoting gut homeostasis and barrier function through diverse mechanisms^10, 125, 194–197, 208, 209^. It is uniquely adapted to colonize the mucus layer and has a diverse enzyme repertoire to degrade and utilize mucin for growth^210^. It also liberates nutrients from mucin to foster the growth of butyrate producers and its production of acetate enhances butyrate production by syntropy^120, 123, 211–214^. In part through its production of acetate, *A. muciniphila*, also stimulates mucin production and does so even better than *Lactobacillus plantarum*, one of the probiotic species previously shown to be protective against morphine dysbiosis and tolerance^215^. Even though pre-morphine microbiota composition was not predictive of non-tolerance, *A. muciniphila* was identified as one of the few biomarkers of non-tolerance prior to morphine (along with *Coprococcus*) and during mid-morphine (Figure 4B) when tolerant mice displayed microbiota instability, and it was notably depleted in the four mice with the lowest ending antinociception (Figure 4C, Supplemental Figure 6, Supplemental Figure 7). One could envision that the timing of its higher abundance could prime the mucosa and microbiota to resist some aspects of dysbiosis and did so better than native *Lactobacillus* and *Bifidobacterium,* which were elevated in tolerant mice pre-morphine (Figure 4B). In accordance with this, vancomycin pre-treatment, which substantially enriches for *A. muciniphila*^216, 217^, curtails tolerance *in vivo* in mice and *in vitro* in neurons even without decreasing microbiota abundance-an affect suggested to be due to the concurrent general decrease in gram positive bacteria, many of which are mutualistic and produce butyrate^37, 116, 201^. Incidentally, vancomycin also improves aspects of autism spectrum disorder, a condition that correlates with low abundance of *A. muciniphila* ^218^, and with low SCFA production^219^. Combined, these suggest the possibility that *A. muciniphila* enhances the accumulation of neuroprotective compounds that curtail tolerance, rather than vancomycin simply eliminating gram positive bacteria and stemming inflammation. Many of the pathobionts identified here and in other studies are gram negative. The abundance of *A. muciniphila* increased in response to morphine as recently shown by others^48^, and even did so modestly in mice that became tolerant, and as such it was not predictive for non-tolerance until post-morphine, where it was substantially depleted in tolerant mice (Figure 4B, Figure 5 and Table 2). As a mucin-degrading bacterium, one could foresee that a higher abundance at the wrong time, or in the absence of butyrate producers that support its immunomodulatory actions, could even compromise barrier functions under morphine stress^48, 220^. This may explain why a probiotic bacterium with so many potential benefits also positively correlates with some neurodegenerative disorders, underscoring the need for caution in employing live probiotics therapeutically while questions persist about modes of action and interaction among the complex microbiota^150, 221–223^.

In contrast with the convergence of microbiota and functions among non-tolerant mice, mice that became tolerant had fewer distinguishing microbiota signatures in common other than an increase in some pathobionts that were also predictive of tolerance (Figure 3, Figure 4B). What was apparent was a subtle difference in the timing of shared patterns of earlier disturbance, illustrated by changes in alpha diversity (Figure 4A), and emergence of taxa included in predictive models during early morphine compared to non-tolerant mice. The common microbiota signatures altered by morphine in tolerant mice include recognized pathobionts linked to intestinal inflammation, including gram negative *Desulfovibrio* and *Bilophila,* and gram positive *Erysipelotrichaceae* and *Rhodococcus,*^224–234^ as well as mucosa-associated taxa, even ones that produce butyrate, (e.g. *Anaerotruncus* and *Butyricicoccus;* ^235^). The association of pathobionts with tolerance is compatible with the model that inappropriate pruning of mutualists, combined with a lack of adaptive responses trained on these rarer pathobionts leads to overgrowth and eventual translocation ^38, 40, 236–239^. Notably, mice that did not develop tolerance also had an increase in pathobiotic taxa and some were even predictive of non-tolerance at some phase of the paradigm (e.g. *Alphaproteobacteria RF32;* Figure 5), underscoring the importance of the temporality of dysbiosis that resulted from a loss of critical stabilizing community members as a contributing to differences in tolerance. The weaker resolution of microbiome associations with tolerance may reflect the fact that tolerant mice exhibited gradients in final antinociception: these mice were less alike than non-tolerant mice (Figure 5 and Supplemental Figure 9). If indeed there are multiple mechanisms of tolerance that contribute to this variability, elucidating microbiome signatures for tolerance would be further constrained by our small cohort size (n=16 mice) where identification of signatures of non-tolerance benefitted from the higher statistical power of categorical assignment. Differentiation of mice that developed tolerance by different mechanisms would further be aided by assessment of additional morphine-driven behaviors that relate to OUD (e.g. mechanical antinociception, dependence, and impaired neuronal plasticity)^240, 241^. More comprehensive behavioral data would support the use of multivariate analyses^242^ to extract meaningful signal in the microbiota of tolerant mice.

## CONCLUSION

Opioids are critical tools in modern medicine and the mainstay for severe and post-operative pain treatment. Unfortunately, opioid use and the ensuing development of tolerance to the analgesic benefits without an accompanying development of tolerance to the respiratory suppressive effects of opioids increases the risk of overdose death as drug dose is necessarily increased to sustain pain management. Only a subset of humans, and as we show here mice, using opioids develop aspects of OUD yet there has been little mechanistic insight to explain this variability. In this study we demonstrate that the gut microbiota contributes to variability in the development of antinociceptive tolerance. Importantly, while we show that microbiota dysbiosis is an inevitable outcome of morphine use, tolerance is not, even in genetically inbred mice from the same vivarium. This emphasizes that, as with humans, mice used to model human disease conditions are not monolithic in part due to the flexible genome conferred by the microbiome. By capitalizing upon this variability, we could ascertain meaningful associations of the microbiota with variability in tolerance and uncouple these associations from hyperalgesia and inflammation. We identify microbial biomarkers of protection from tolerance and show how functions provided by these microbes in the community, not individual microbes *per se,* underlie protection. We identified the function of butyrate production, and by proof-of-concept through dietetic supplementation, demonstrated that it can reduce tolerance. Due to the foundational impact of germ theory of disease on the discipline of microbiology, microbes have been historically categorized as the problem, and this pervasive view has impeded progress in understanding the contributions of mutualistic community members to complex health conditions. Our study reinforces the importance of having a more holistic view to understand interactions between the microbiota and host. In the case of OUD, a better understanding of interactions among microbiota and between host and microbiota during morphine use could reveal underlying local or systemic mechanisms of tolerance, an understanding of which could reveal therapeutic targets and produce opportunities for effective pain management while limiting dangerous opioid side effects. Taken together, our studies indicate that the gut microbiota could be leveraged to reduce tolerance to the analgesic benefits of opioids and thereby reduced the risk of OUD.

## ACKNOWLEDGMENTS

We thank Sarah W. Gooding, Madeline King, Joshua Gipoor, and Benjamin Wasson for assistance with data curation and collection; Maria Emanuel, Stephen Fiering, and Stephen Jones for their immeasurable support. We are grateful for the expertise of the staff at the Hubbard Center for Genome Studies at UNH and especially W. Kelley Thomas for advice, and Anthony Westbrook for bioinformatics assistance. We are grateful for the deep knowledge, helpful guidance, assistance and patience of Linnea Morley and Dean Elder at the Animal Resource Office at UNH.

## DATA AVAILABILITY STATEMENT

The 16S amplicons that support the findings of this study are available in NCBI under the accession number PRJNA1098090. The authors confirm that all processed data supporting the findings of this study are available within the article and/or its supplementary materials.

