## Supplemental Figures for "Gut dysbiosis was inevitable, but tolerance was not: temporal responses of the murine microbiota that maintain its capacity for butyrate production correlate with sustained antinociception to chronic morphine"

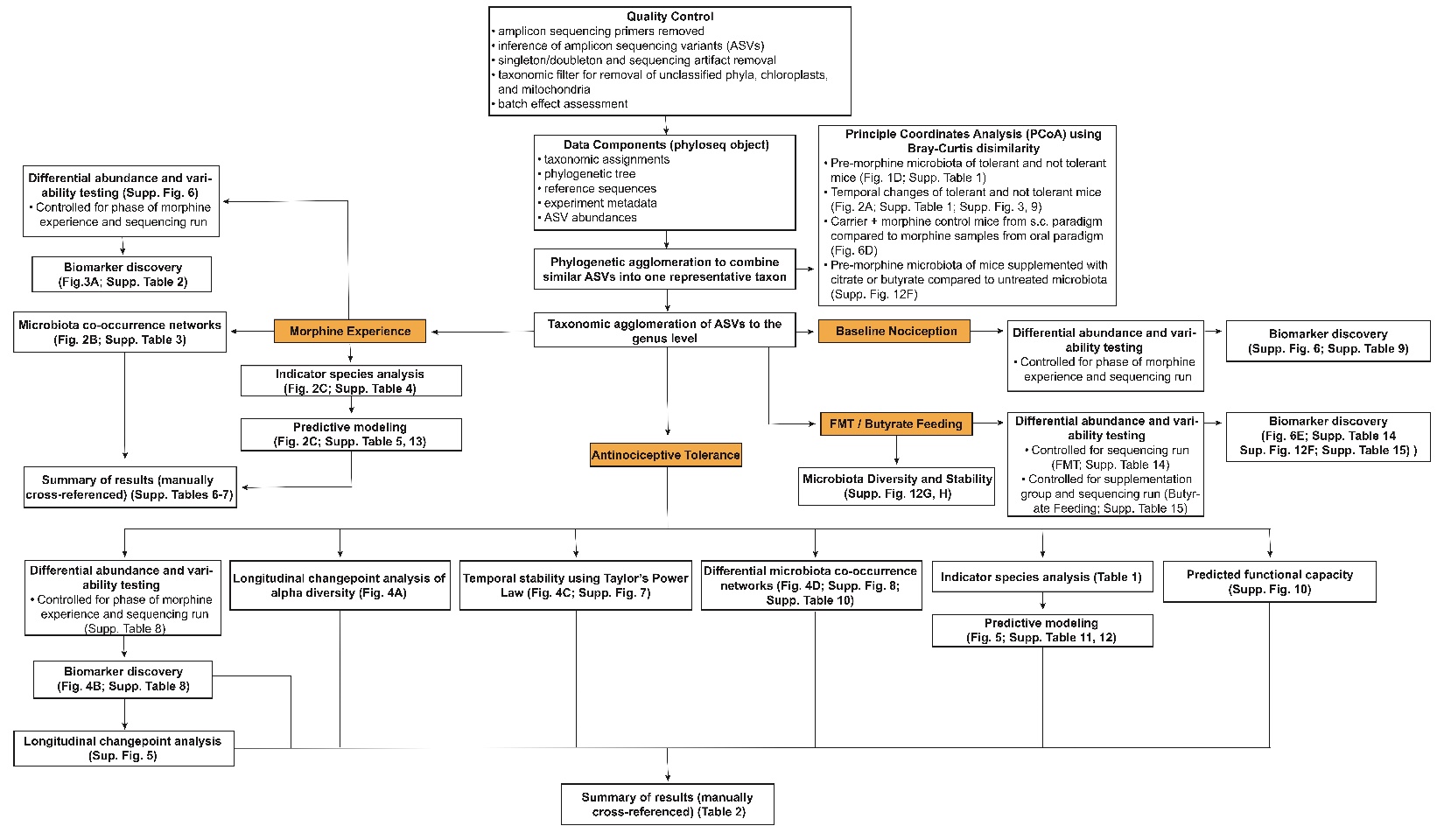
**Supplemental Figure 1: Pipeline of 16S rDNA sequencing read processing and subsequent analysis of amplicon sequence variants (ASVs) for associations with morphine experience, antinociceptive tolerance and baseline nociception.** Demultiplexed raw sequencing reads were processed using cutadapt, DADA2, and custom scripts for post-processing artifact removal to generate a quality-controlled dataset. Taxonomy was assigned using the GreenGenes reference database (v13.8) (Second Genome Inc). All data components (i.e., taxonomic assignments, experiment metadata, ASV counts, ASV sequences, and a phylogenetic tree) were stored together using the phyloseq package for downstream analyses. Subsequent analyses utilized ASV datasets that were either agglomerated by phylogeny (similar ASVs into one representative taxon) or taxonomy (genus level) to investigate microbiome covariates with morphine experience, nociceptive tolerance, baseline nociception, fecal microbiota transplantation, and citrate or butyrate supplementation. All analyses and visuals were produced using R Studio (v4.2.0).

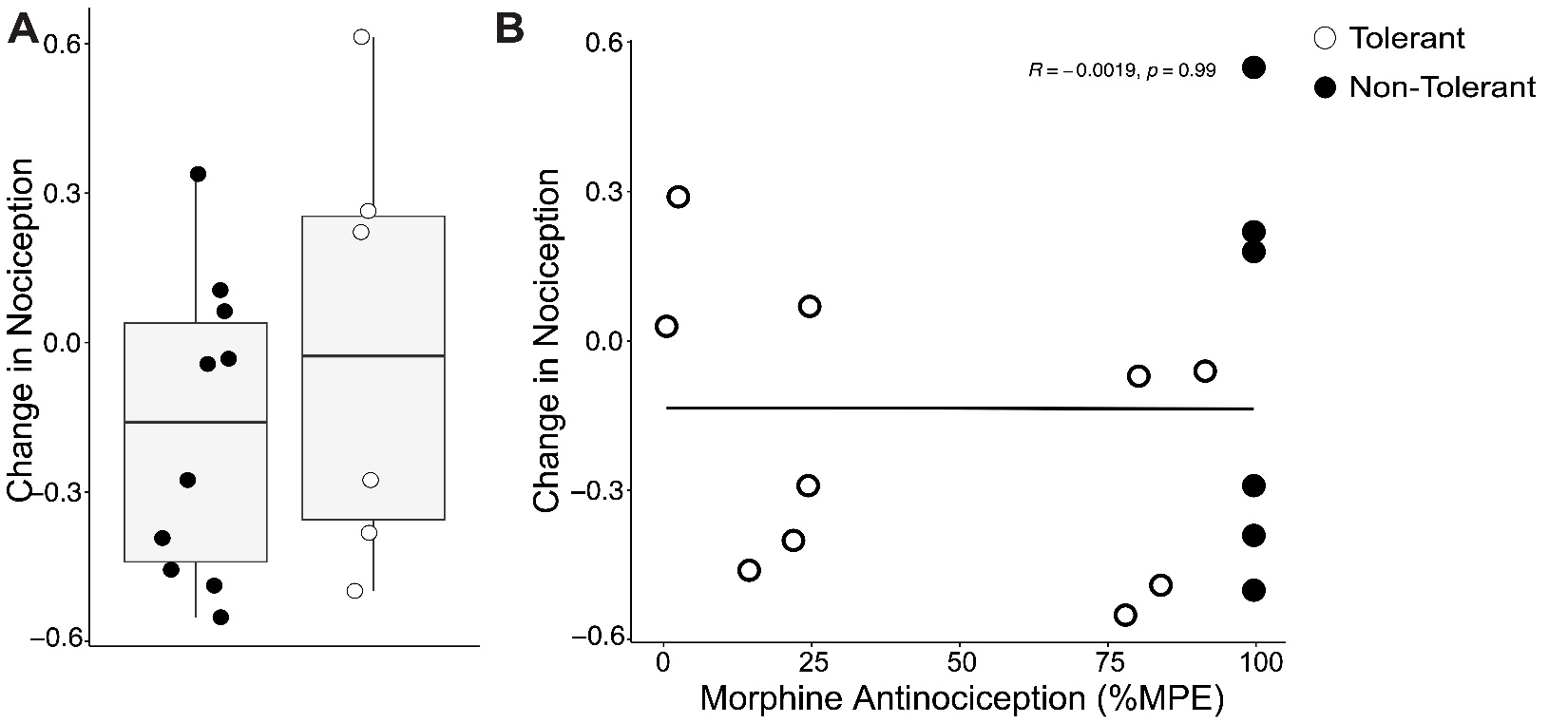
**Supplemental Figure 2: Mice developed minimal hyperalgesia which did not correlate with tolerance.** A) Change in nociception was calculated using the following formula: Day 1 baseline latency (s) - Day 127 baseline latency (s). B) Linear regression of change in nociception with degree of tolerance (y = -0.13 - 0.000016x; R2 = -0.0019; Pearson’s correlation p-value = 0.99).

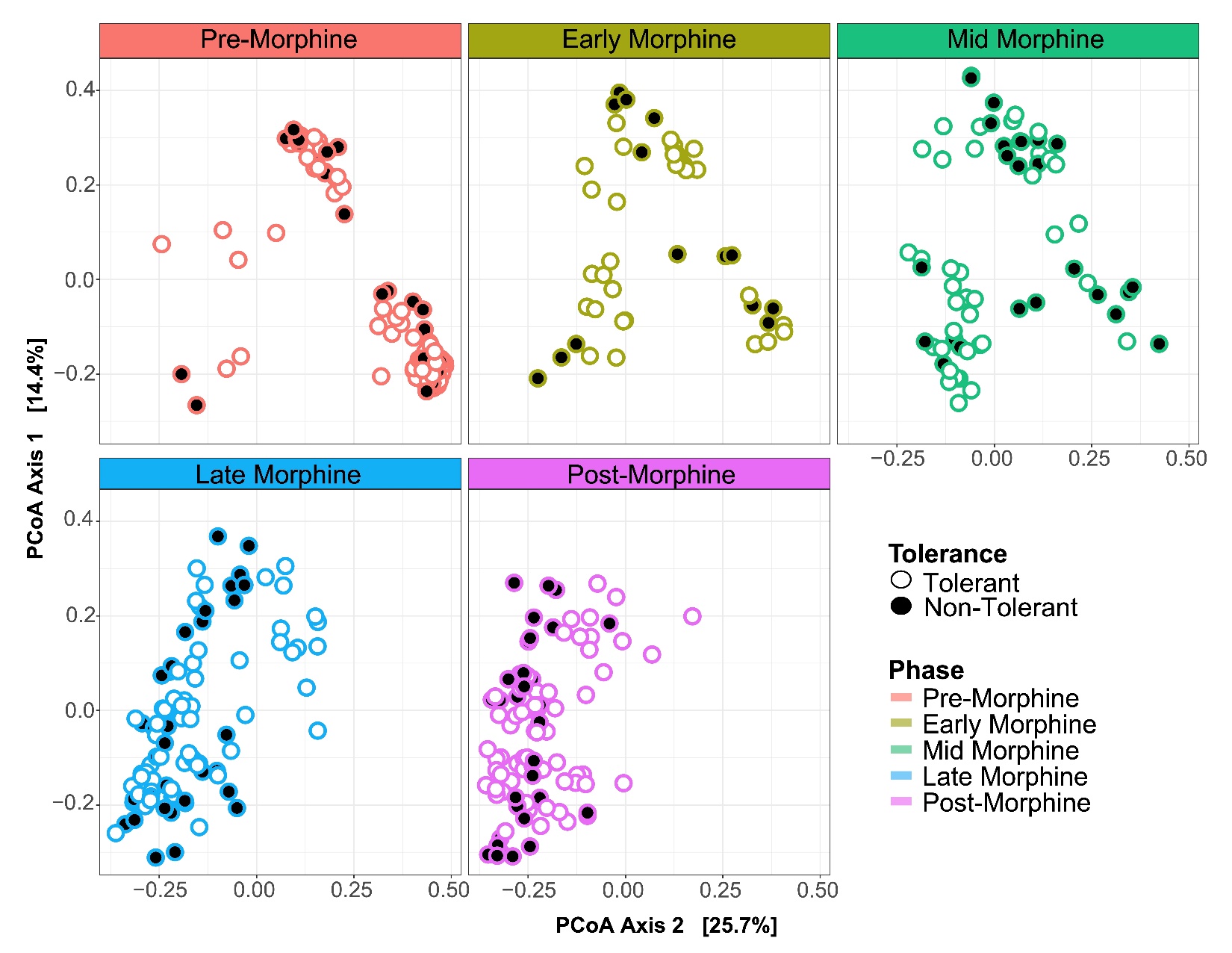
**Supplemental Figure 3: Microbiomes of tolerant and non-tolerant mice do not form separate clusters at any phase of morphine exposure exemplifying how microbiota associations with tolerance are latent variables.** Temporal changes in the gut microbiota β-diversity of mice that were tolerant or non-tolerant (All ASVs, Bray-Curtis dissimilarity) as visualized by principal coordinates analysis (PCoA).

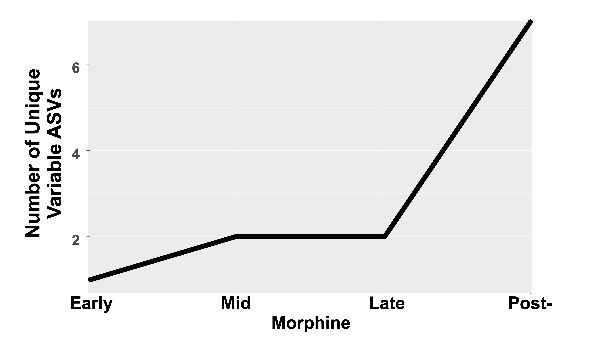

**Supplemental Figure 4: Morphine-driven increases in unique differentially variable ASVs.** The number of community members that were differentially variable due to morphine consumption through time were identified using corncob Wald-type chi-squared test ^88^ (See Supplemental Table 2 for details).

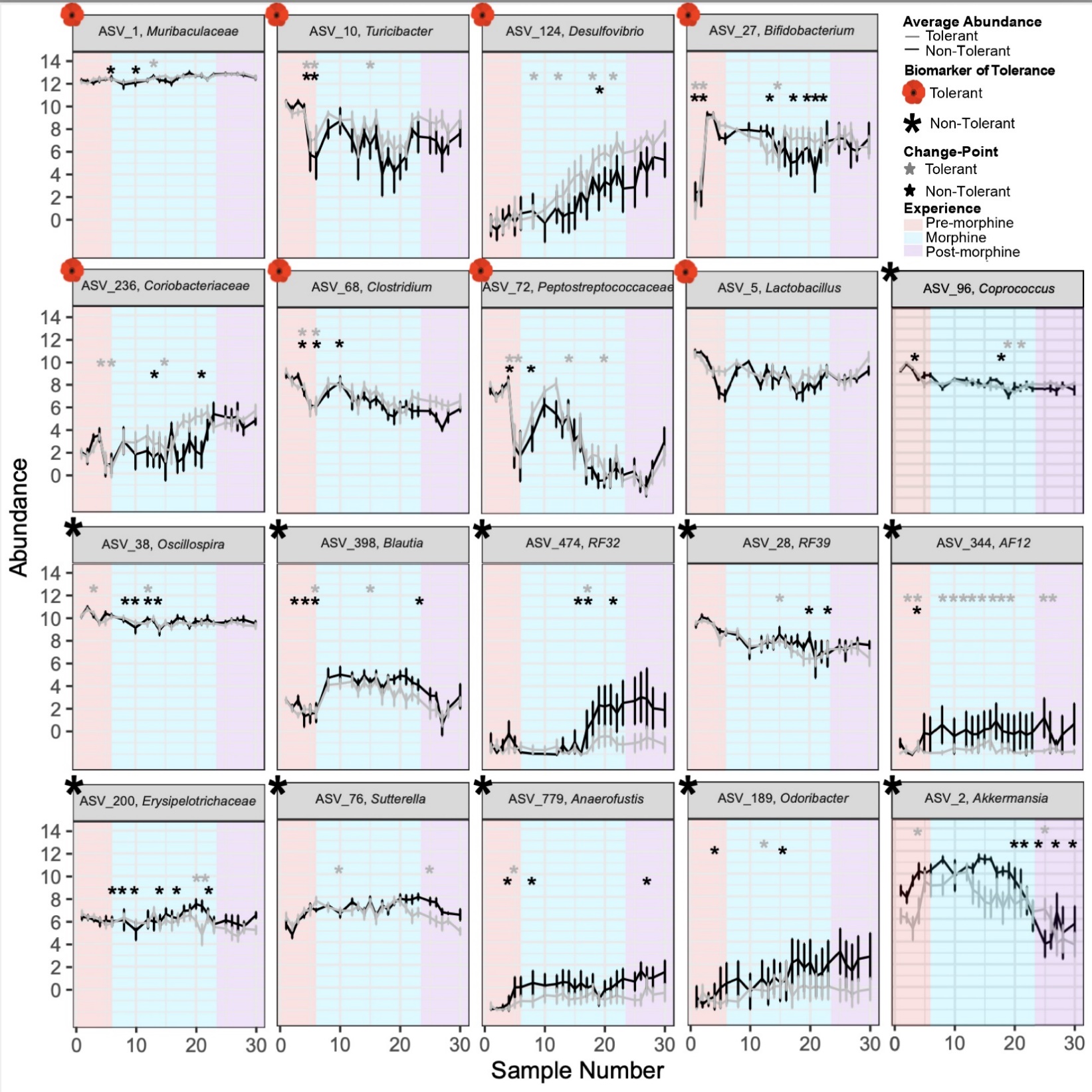

**Supplemental Figure 5: Comparisons and estimated points of significant change in the longitudinal distribution of biomarkers of tolerance and non-tolerance exemplify instability in tolerant mice.** Significant temporal change points were detected using Bayesian change-point analysis of the centered log-ratio transformed abundances of biomarkers of tolerant (grey line) or non-tolerant (black line) mice identified at one or more than one point in the paradigm or when all experiences were combined (as described in Figure 4B) and plotted temporally by fecal sample number. Significant change points identified by asterisk. Error bars represent standard error.

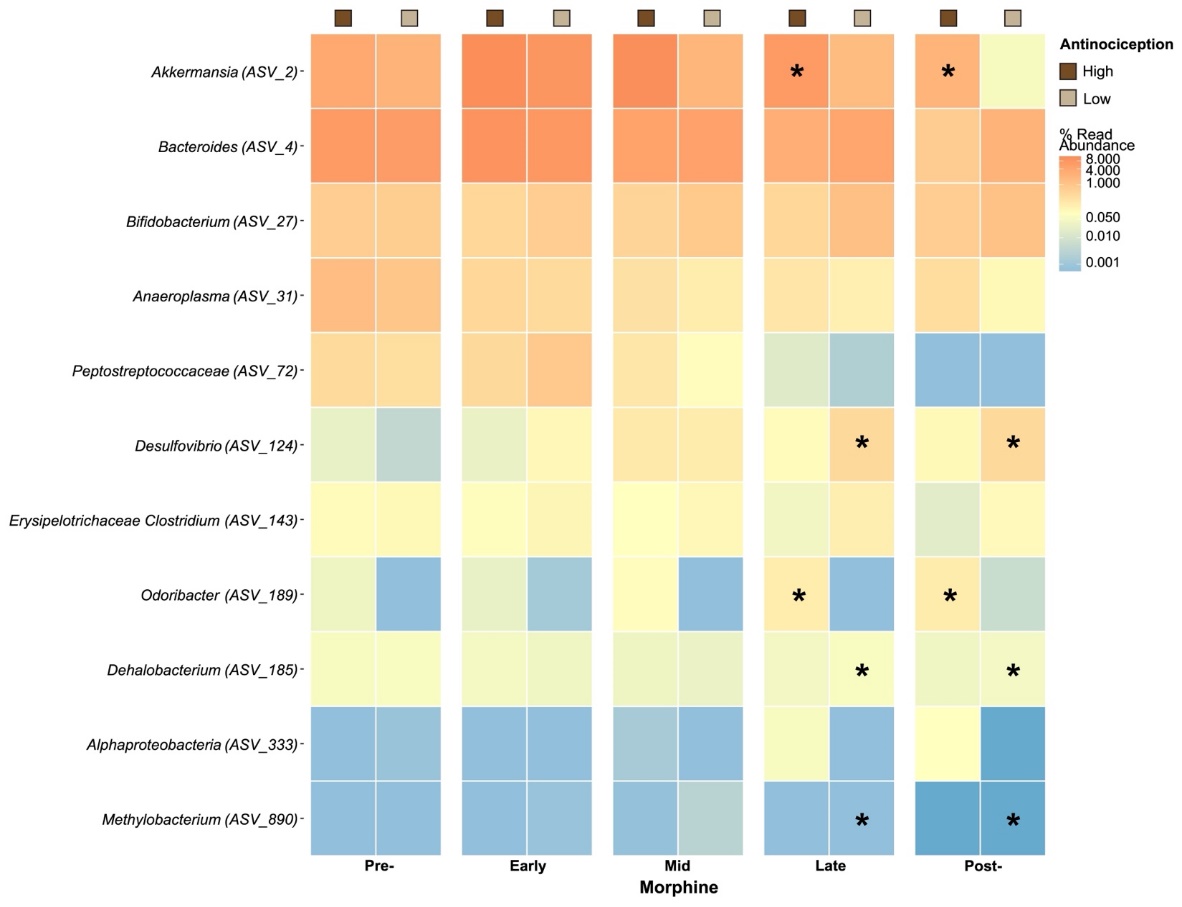

**Supplemental Figure 6: The gut microbiota of mice with low baseline morphine antinociception (<100%) differed from those with 100% antinociception.** Biomarkers of low baseline morphine antinociception were identified from among the community members whose abundance was significantly different (as identified using corncob regression models)^88^, followed by a linear discrimination analysis (LDA) of effect size (LEfSe)^89^ to identify genera that most likely explain differences in morphine efficiency (asterisk). Community members are labeled at the lowest taxonomic classification available. See Supplemental Table 9 for details.

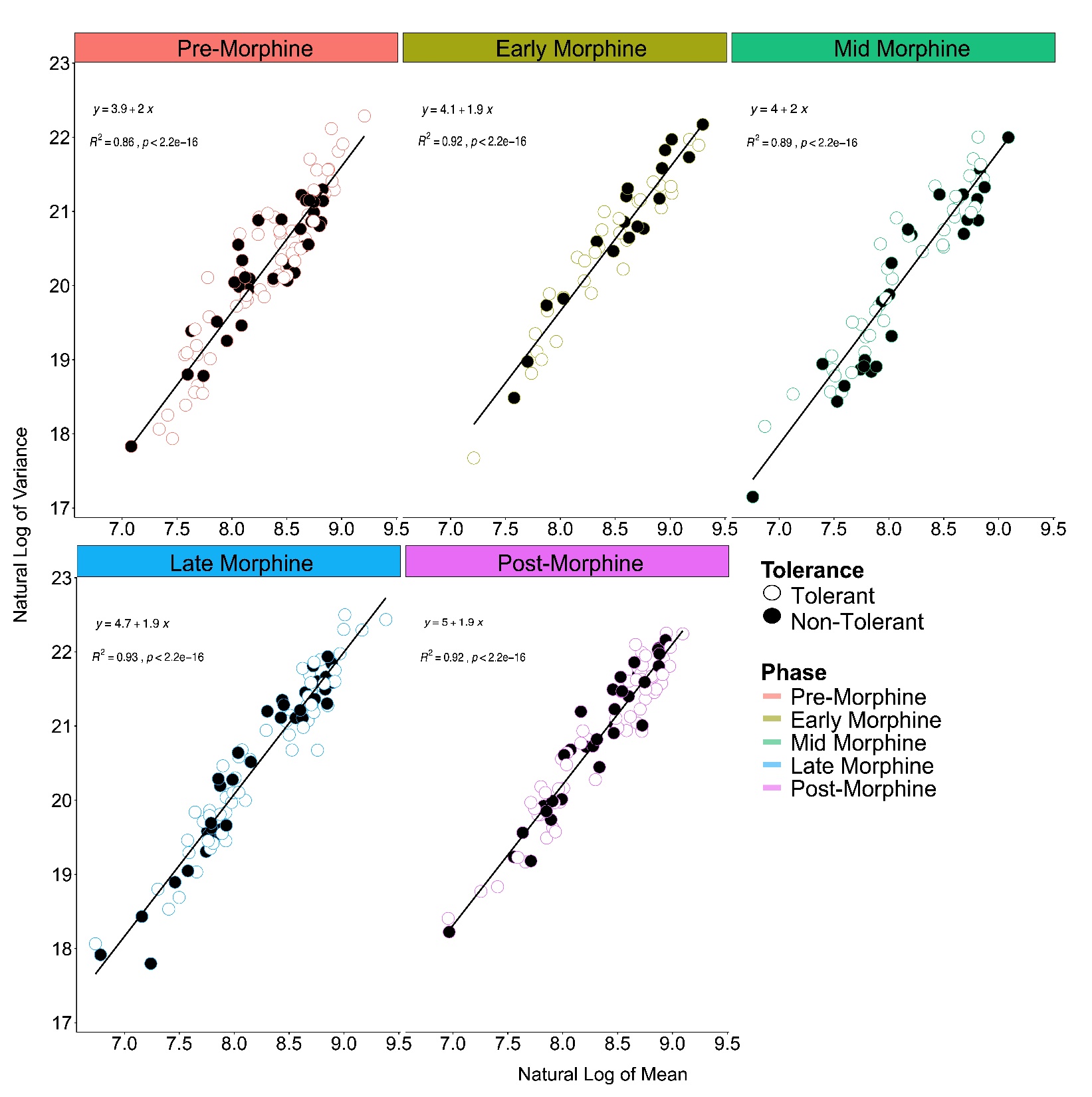

**Supplemental Figure 7: All microbial communities exhibit normal stochasticity independent of phase of morphine experience or development of tolerance.** Type I Taylor’s Power Law (mean-variance relationship of all microbial communities over time) for microbiota of tolerant and non-tolerant mice across all phases of the morphine experience. Pearson’s correlation determined whether there was significant aggregation of microbiota at each phase guided by a previously published threshold (R^2^ above 0.34) ^91^. See methods for details.

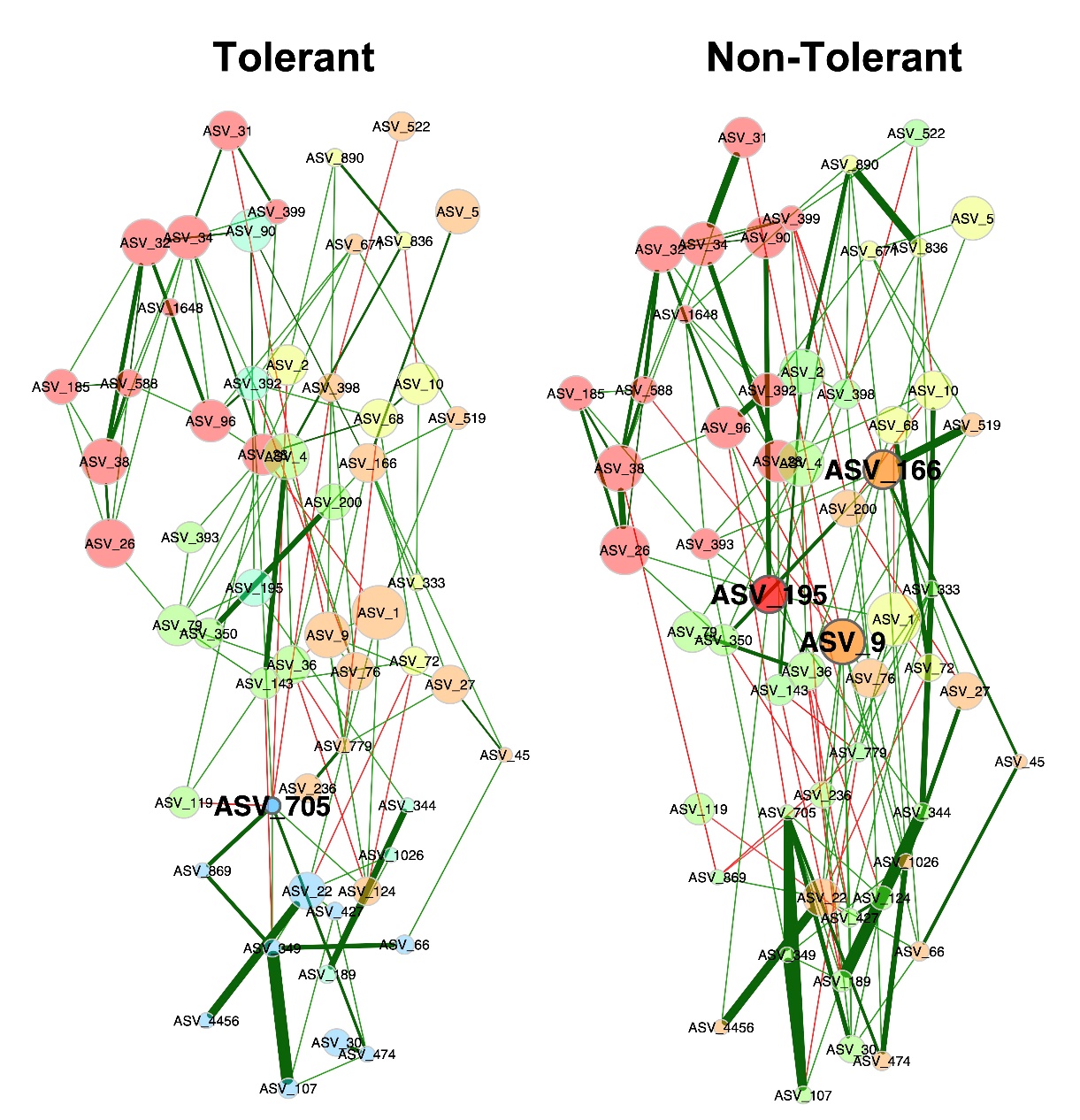

**Supplemental Figure 8:** **Global network properties of tolerant and non-tolerant mice did not differ***.* Comparison of bacterial association networks of non-tolerant (left) and tolerant mice (right) using the SPRING method in NetCoMi^94^. Edges are colored by sign (positive: green, negative: red), representing estimated associations between community members. *Adlercreutzia* (ASV_166), *Mogibacteraceae* (ASV_195), *Allobaculum* (ASV_9) and *Peptococcaceae* (ASV_705) are the community members identified as hubs (bolded text and borders). Eigenvector centrality was used for scaling node sizes and colors and representing clusters of genera (determined using greedy modularity optimization). Clusters have the same color in both networks if they share at least two taxa, clustersbetween tolerant and non-tolerant mice are not similar (ARI 0.275, p-value=0) and specific associations defining these differences are presented in Figure 4D. Nodes that are not connected in both groups were excluded.

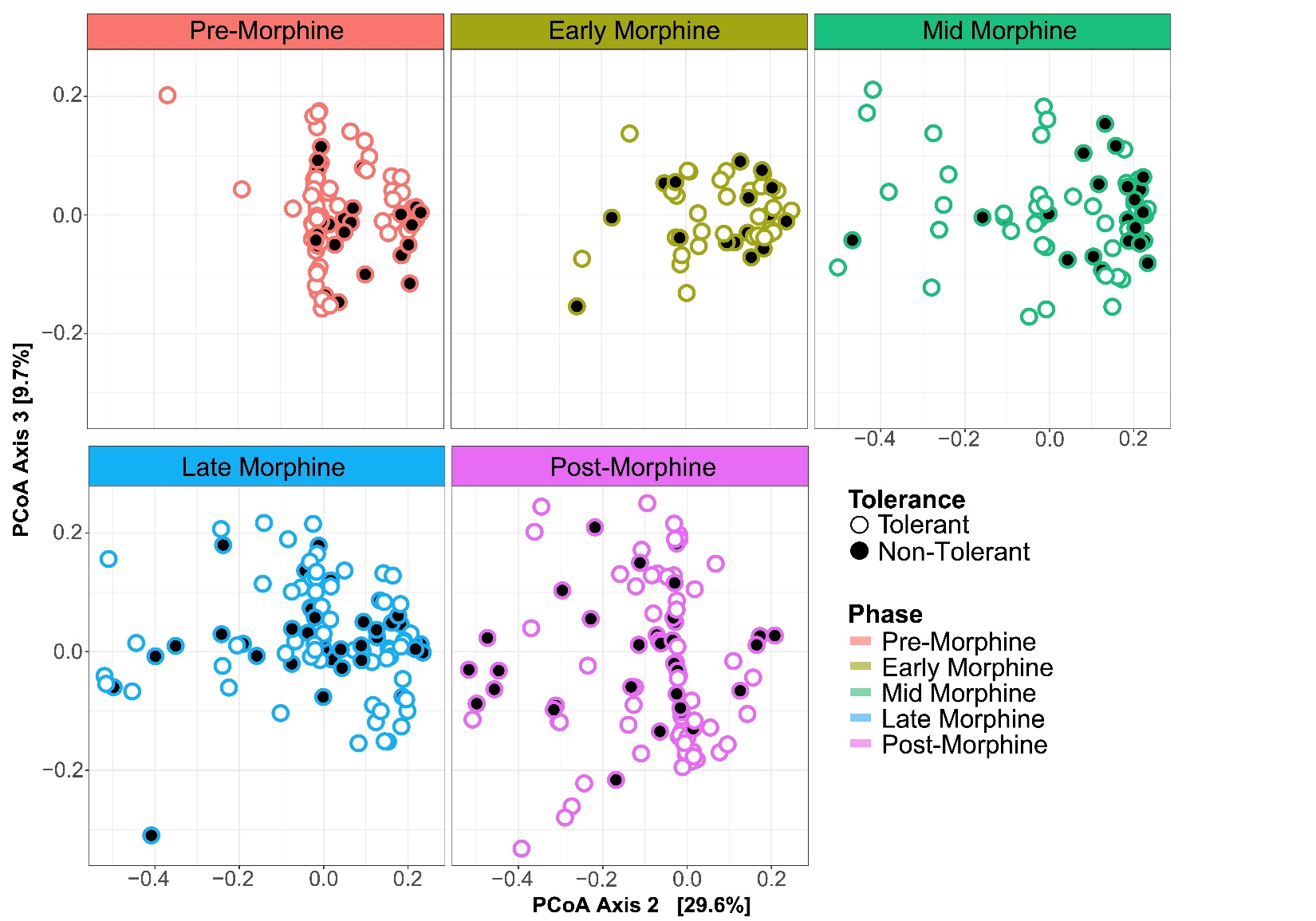

**Supplemental Figure 9: Temporal changes in microbiota β-diversity of mice using only biomarker genera exemplify more variability among tolerant mice mid-morphine.** Differentially abundant taxa from Figure 4B, weighted UniFrac distance, were visualized by principal coordinates analysis (PCoA). . Microbiota from non-tolerant mice cluster mid-morphine whereas those from tolerant mice vary more from each other, especially during mid-morphine. These relationships were not visible on Axis 1 that explained 33.4% of the microbiome variability.

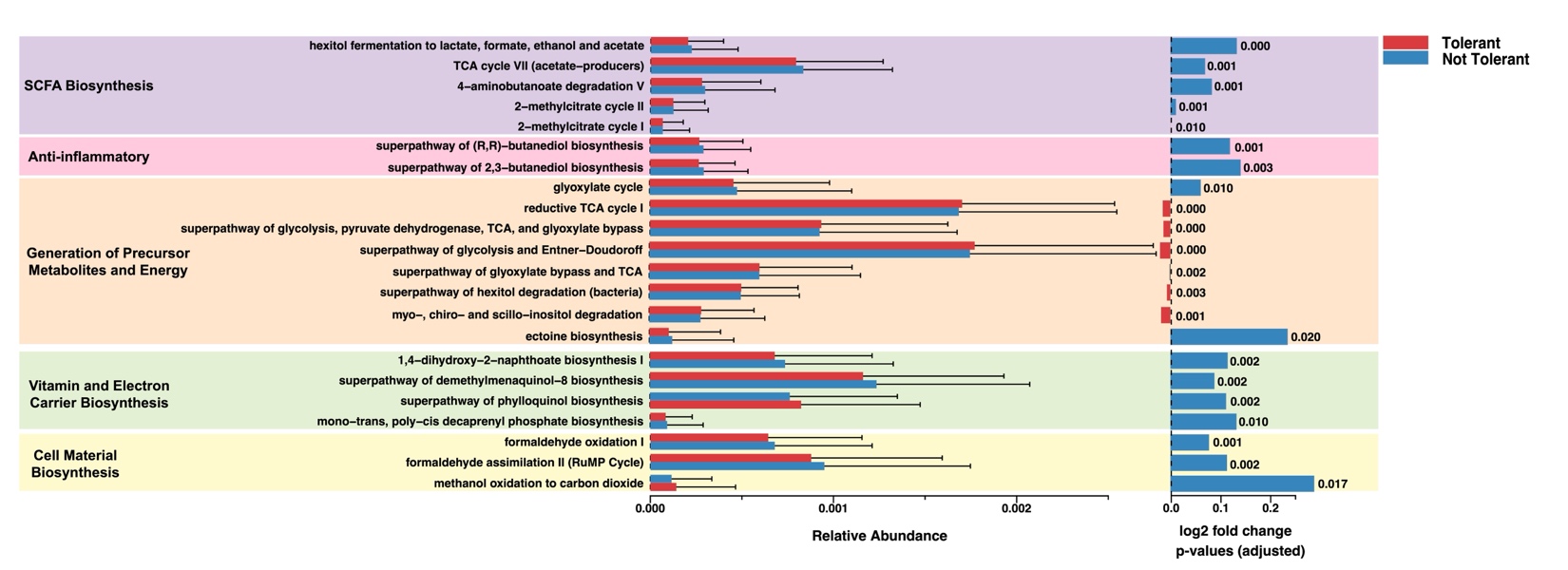
**Supplemental Figure 10: Functional capacity of the gut microbiota differs between tolerant and non-tolerant mice.** Predicted functional profiles (metabolic pathways and abundances) derived from 16S rRNA sequences were generated using PICRUSt2^95^. Differential abundance testing of predicted functional pathways between tolerant (red) and non-tolerant mice (blue) was compared across three models, including ALDEx2^102^, DESeq2^103^, and edgeR^104^ using the ggpicrust2^101^ package (v1.7.3) in RStudio (v4.2.0). Only significant predicted functional pathways using the three differential abundance tests are shown.

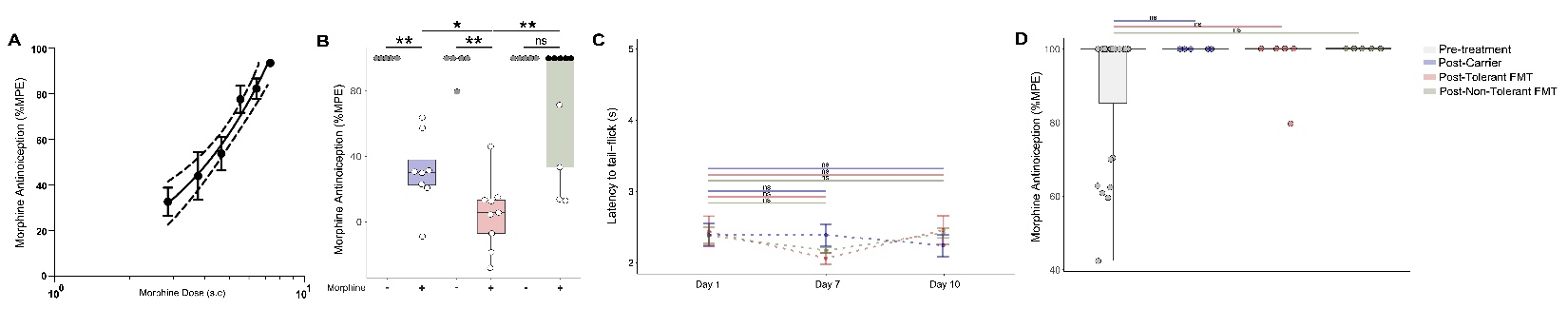
**Supplemental Figure 11: Influence of FMT on antinociception and nociception .** A) Linear regression of dose response curves (3, 4, 5, 3+3, 5+2 and 8 mg/kg) used to assign the potency of 5 mg/kg (ED_60_) and 5+2 mg/kg (ED_90_) morphine (m.s.). Error bars are SEM and outside dashed margins are 95% CI of the linear regression. B) Day 10 antinociception (5 + 2 mg/kg m.s. s.c.) using the radiant heat tail-flick assay and reported as % MPE. Kruskal-Wallis test followed by pairwise Wilcox test for multiple comparisons determined significant differences between m.s.-treated carrier and FMT groups and between m.s. treatments and their saline controls (ns p>0.05; *p<0.05; **p<0.01). C) Baseline latencies to tail-flick where Kruskal-Wallis test followed by pairwise Wilcox test for multiple comparisons on day 1, 7, and 10 where a decrease in latency would reflect hyperalgesia. Scale is the same as Figure 6F. Error bars are SEM. D) Morphine antinociception (5+2 mg/kg s.c. m.s.) as determined on day 10 using the radiant heat tail-flick assay for carrier and FMT groups that did not experience morphine during days 1-9 compared to antinociception of mice pre-treatment . Data is reported as maximum possible effect (% MPE). Kruskal-Wallis test followed by pairwise Wilcox test for multiple comparisons were used to determine whether FMT treatments alone altered morphine antinociception.

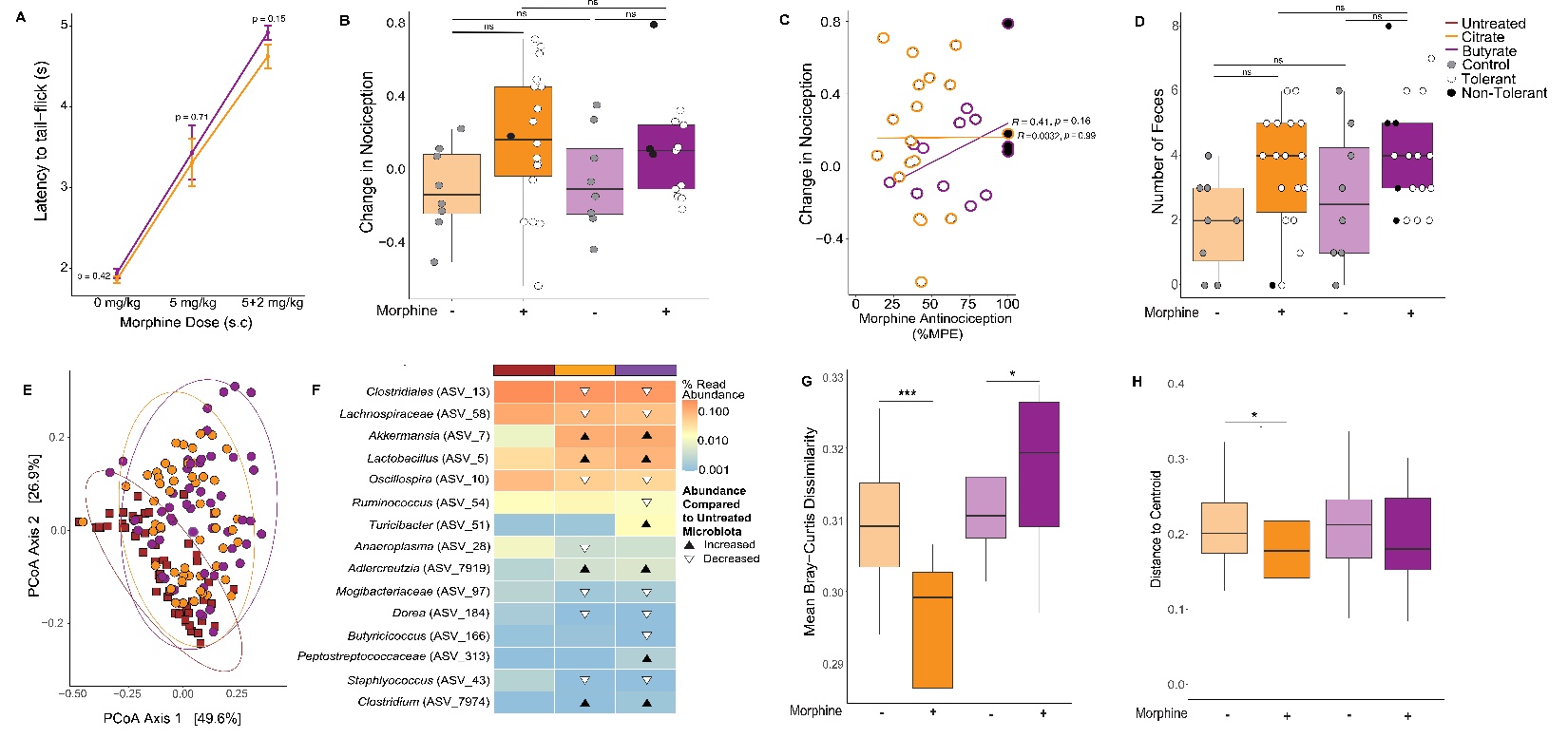
**Supplemental Figure 12: Influence of citrate and butyrate supplementation on antinociception latencies, change in nociception, defecation, and gut microbiota.** A) Day 1 latencies to tail flick to radiant heat where Kruskal-Wallis test followed by pairwise Wilcox test for multiple comparisons were used to determine whether citrate and butyrate supplementation alone altered nociception and antinociception . Error bars are SEM. B) Change in nociception (day 1 latency - day 10 latency S) was calculated for each mouse ANOVA test followed by Tukey’s HSD test for multiple comparisons was used to determine whether citrate or butyrate treatment altered nociception . A positive change in baseline nociception would reflect hyperalgesia. Five butyrate mice were excluded from nociceptive analyses due to technical difficulties on day 1 during their baseline assessments. C) Linear regression of change in nociception with degree of tolerance after 10 days of morphine treatment in citrate (y = 0.16 + 0.000061x; R2 = 0.0032; Pearson’s correlation p-value = 0.99) or butyrate groups (y = -0.2 + 0.0043x; R2 = 0.41; Pearson’s correlation p-value = 0.16). D) Reduced gut transit from chronic morphine was quantified by number of fecal pellets produced following reversal with the MOR antagonist naloxone. ANOVA followed by Tukey’s HSD test for multiple comparisons evaluated whether morphine caused significant constipation and whether thiw was influenced by butyrate supplementation. . E) Shifts in gut microbiota β-diversity (genus level, Bray-Curtis dissimilarity) in response to dietic supplementation with citrate or butyrate compared to microbiota in untreated animals as visualized by principal coordinates analyses (PCoA). F) Biomarkers of citrate or butyrate supplementation were identified from among the community members whose abundance was significantly altered by citrate or butyrate using corncob binomial regression models ^88^, followed by a linear discrimination analysis (LDA) of effect size (LEfSe) ^89^. Black and white triangles indicate whether the abundance of a community member increased (black triangle) or decreased (white triangle) compared to untreated microbiota. Community members are labeled at the lowest taxonomic classification available. See Supplemental Table 15 for details. Comparisons of gut microbiota (G) β-diversity (genus level, mean Bray-Curtis dissimilarity) and (H) variability (genus level, Bray-Curtis dissimilarity) between the microbiota of morphine-treated citrate and butyrate supplementation groups and their pre-morphine microbiota determined whether butyrate supplementation impacted community composition and stability. Pairwise (G) PERMANOVA and (H) permutation tests determinedsignificant differences between the microbiota of morphine-treated citrate and butyrate supplementation groups and their pre-morphine microbiota (*p<0.05;***p<0.001).
